# Reduction of embryonic *E93* expression as a key factor for the evolution of insect metamorphosis

**DOI:** 10.1101/2022.10.04.510826

**Authors:** Ana Fernandez-Nicolas, Gabriela Machaj, Alba Ventos-Alfonso, Viviana Pagone, Toshinori Minemura, Takahiro Ohde, Takaaki Daimon, Guillem Ylla, Xavier Belles

## Abstract

The early embryo of the cockroach *Blattella germanica* exhibits high *E93* expression. In general, E93 triggers adult morphogenesis during postembryonic development, but in the cockroach E93 is also crucial in early embryogenesis. Moreover, the embryonic levels of *E93* expression are high in hemimetabolan insects, while in holometabolans they are very low. They are also low in Thysanoptera and in Hemiptera Sternorrhyncha with postembryonic quiescent stages, as well as in Odonata, the nymph of which is very different from the adult. In ametabolans, such as the Zygentoma *Thermobia domestica, E93* expression levels are very high in the early embryo, whereas during postembryonic development they are medium and relatively constant. Given that embryogenesis of hemimetabolans yields an adultiform nymph, we speculate that E93 plays some sort of adult triggering role in the embryo of these species. We conjecture that the reduction of *E93* transcript levels in the embryo has been instrumental in the evolution of insect metamorphosis. The suppression of *E93* expression during the nymphal period, and its concentration in the preadult stage, is consubstantial with the emergence of hemimetaboly. As such, attenuation of *E93* expression in the embryo could have resulted in a larval genetic program and the emergence of holometaboly. Independent decreases of *E93* expression in the embryo of Odonata, Thysanoptera, and different groups of Hemiptera Sternorrhyncha would have allowed the development of modified juvenile stages adapted to specific ecophysiological conditions.

## Introduction

The young William Harvey was a hot-tempered man who often played with a dagger which he wore at his waist (1). It is not surprising, then, that he felt particularly fond of the study of blood circulation. However, his contribution to the study of animal reproduction, including insects, is no less important. Everything comes from the egg, *ex ovo omnia*, is the motto that sums up Harvey’s work (2). Our present contribution contains the message that in the study of insect metamorphosis, despite being a post-embryonic process, it is crucial to include the analysis of embryo development.

Insect metamorphosis is generally classified into two main types — hemimetaboly and holometaboly— while ametaboly refers to non-metamorphosing species. In hemimetabolans, the hatched nymphs already have the adult body structure, and only at the end of the nymphal period does the insect transform into the adult stage, fully winged and sexually competent. In holometabolans, embryo development gives rise to a larva that is morphologically different from the adult. At the end of the larval period, the insect molts to the pupal stage, a kind of intermediate between the larva and the adult, and then to the fully winged and sexually competent adult (3).

The above definition of hemimetaboly and holometaboly means that embryo development determines what kind of juveniles and what type of metamorphosis will follow. However, metamorphosis scholars rarely pay attention to embryogenesis. Although review papers recognize that studying embryogenesis is important to understand metamorphosis (3–6), experimental studies in this regard are scarce (7–10).

A comparative analysis of RNA-seq data for the fly *Drosophila melanogaster* and the cockroach *Blattella germanica* (11) revealed that the gene *E93* is expressed in pupae in *D. melanogaster*, while in *B. germanica* it is expressed in the last nymphal instar, but also at early stages of embryogenesis. E93 triggers adult morphogenesis (12, 13), via a mechanism involving the modulation of chromatin accessibility (14, 15). Thus, we wondered what would be the action of E93 in the embryo of *B. germanica*, and how general would be the occurrence of high levels of embryonic expression in hemimetabolans (and low levels in holometabolans). Ultimately, we speculated that, given that embryogenesis yields an adultiform nymph in hemimetabolans, E93 might play some sort of adult-specifying role in the embryo of these species.

## Results and discussion

### E93 is required for cockroach embryogenesis

Real-time quantitative reverse transcription PCR (qRT-PCR) measurements showed that maximal levels of *E93* transcripts in the embryo of *B. germanica* occur on day 1 of development (ED1) (Fig. 1*A*). These levels subsequently decrease and remain low throughout embryogenesis. To study the role of embryonic E93, we used maternal RNAi. Thus, 5-day-old females (AdD5) were injected with 3 μg of dsRNA targeting *E93* mRNA (dsE93). Control females received the same dose of control dsRNA (dsMock). All females were then allowed to mate. *E93* expression in embryos from dsE93-treated females was significantly lower (71%) than in controls (Fig. 1*B*). All control females (n=40) formed an ootheca on day 8 of the adult stage (AdD8), which contained viable embryos that became hatchlings 18 days later (35–40 emerged nymphs per ootheca). Females treated with dsE93 (n=36) also formed an ootheca on AdD8, and 50% of them gave first instar nymphs 18 days later (between 35 and 40 emerged nymphs per ootheca). However, 27.8% dropped the ootheca between ED2 and ED3 (12–17% embryogenesis), whereas 22.2% transported the ootheca beyond day 18, dropping it between ED19 and ED21. No nymphs emerged from these 18 oothecae.

**Fig. 1.**
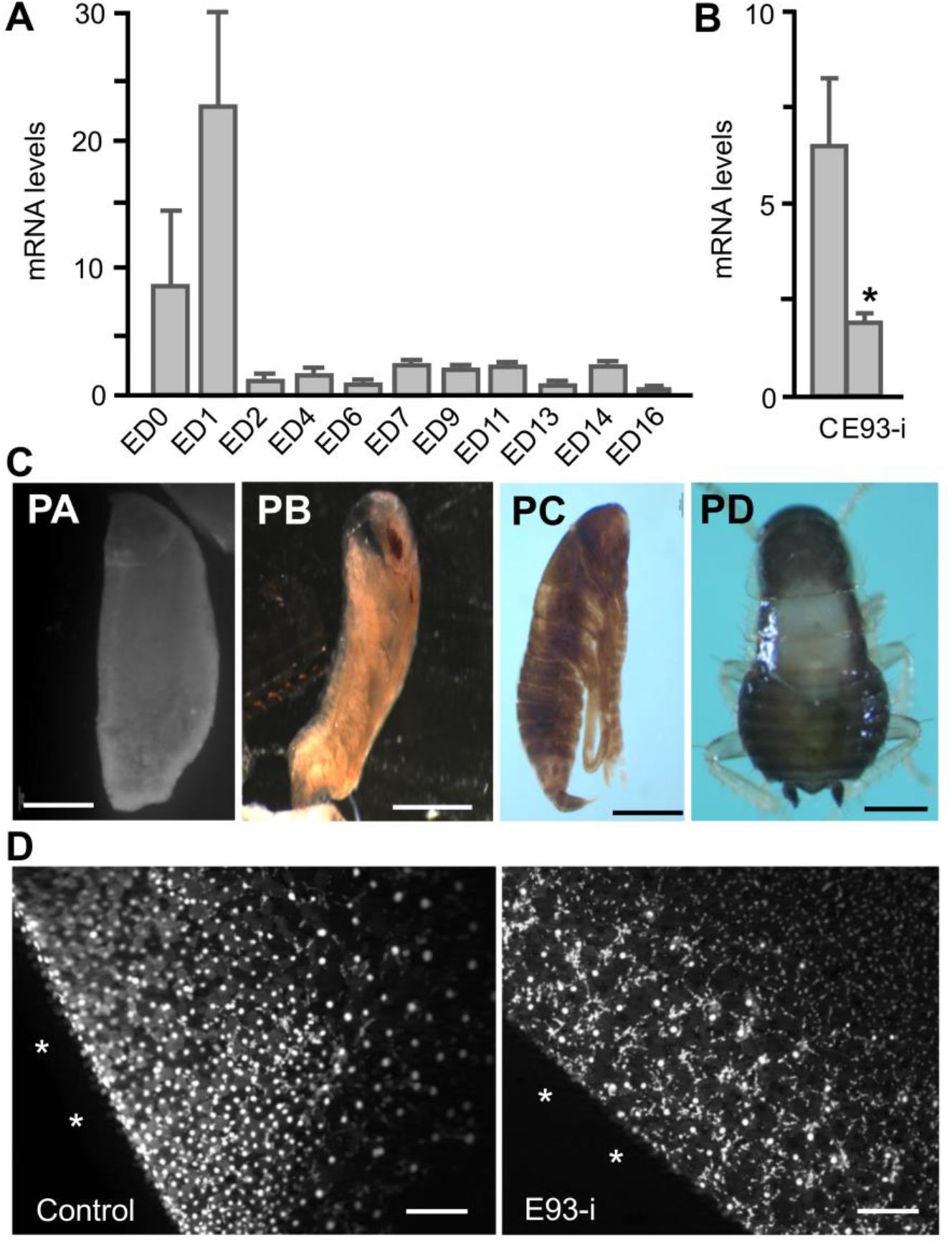
Effects of E93 depletion on embryo development in *Blattella germanica*. (*A*) *E93* expression in embryos of different ages from day 0 (ED0) to day 16 (ED16). (*B*) *E93* expression after maternal RNAi at ED0 in control (C) and treated (E93-i) insects. (*C*) Phenotypes PA (25.5% of cases) resulting from E93 depletion: embryos that interrupted development around the formation of germ-band anlage; PB (14.4% of cases): embryos that interrupted development between Tanaka stages 10 and 15; PC (19.4% of cases): apparently well-formed nymphs, with dark cuticle; and PD (40.6% of cases): embryos which, although they are similar to pre-hatched nymphs, could not hatch, but if the chorion was broken, they looked like naturally eclosed first instar nymphs, as shown in the picture. (*D*) Accumulation of energids in the ventral side of the embryo (white asterisks) in control and E93-i specimens, in ED2. In (*A*) and (*B*), each qRT-PCR expression measurement represents three biological replicates, and results are expressed as copies of the examined transcript per 1000 copies of BgActin-5c mRNA; data are represented as the mean ± SEM. In (*B*), the asterisk indicates statistically significant differences with respect to controls (p-value < 0.05), calculated on the basis of the REST (28). Scale bars in (*C*): 500 μm; in (*D*): 200 μm.

On ED19 we examined 278 embryos from the eight oothecae formed by dsE93-treated females that did not hatch on ED18. Of these, 25.5% presented interrupted development in a pre-blastoderm stage, or stage 1-2 of embryogenesis, as defined by Tanaka (16) (Fig. 1*C* PA). Moreover, 14.4% had interrupted development between Tanaka stages 10 and 15, showing a vermiform aspect, with poorly defined segments and appendages, an imperfectly sealed dorsal closure, reduced abdomen or imperfect eyes (Fig. 1*C*, PB). A total of 19.4% were apparently well-formed nymphs, but with an intensely sclerotized cuticle (Fig. 1*C*, PC), whereas 40.6% were practically indistinguishable from control pre-hatched nymphs (Fig. 1*C*, PD).

We then studied embryos from oothecae from dsE93-treated females that were dropped on ED2, compared with ED2 embryos from control females. We analyzed 350 embryos from the latter (control embryos) and 235 E93-depleted embryos. Control embryos were between Tanaka stages 1 and 2, and showed a high density of energids accumulating on the ventral side of the egg, where they would contribute to formation of the germ-band anlage (Fig. 1*D*, left panel). Although E93-depleted embryos were also around Tanaka stages 1 and 2, the accumulation of energids in the ventral side of the egg was lower than in controls, and their distribution was more heterogeneous, forming discrete groups separated from each other (Fig. 1*D*, right panel).

### E93 depletion results in important transcriptomic changes in the early embryo

We knocked down *E93* expression in early embryos using maternal RNAi, and prepared RNA libraries of controls and E93-depleted samples at ED2, in other words one day after the *E93* expression peak. We performed the E93 knockdown experiment in two independent batches, and sequenced three libraries from each batch of E93*-*depleted embryos and their respective controls (SI Appendix, Table S1). We obtained a minimum of 60 million reads for each sample, 90% of which were mapped to the reference genome (SI Appendix, Table S2 and Dataset S1). Principal component analysis (PCA) and batch correction analyses (Fig. 2*A* and SI Appendix, Fig. S1) resulted in six control and five treated bona fide libraries, which were used for differential gene expression studies.

**Fig. 2.**
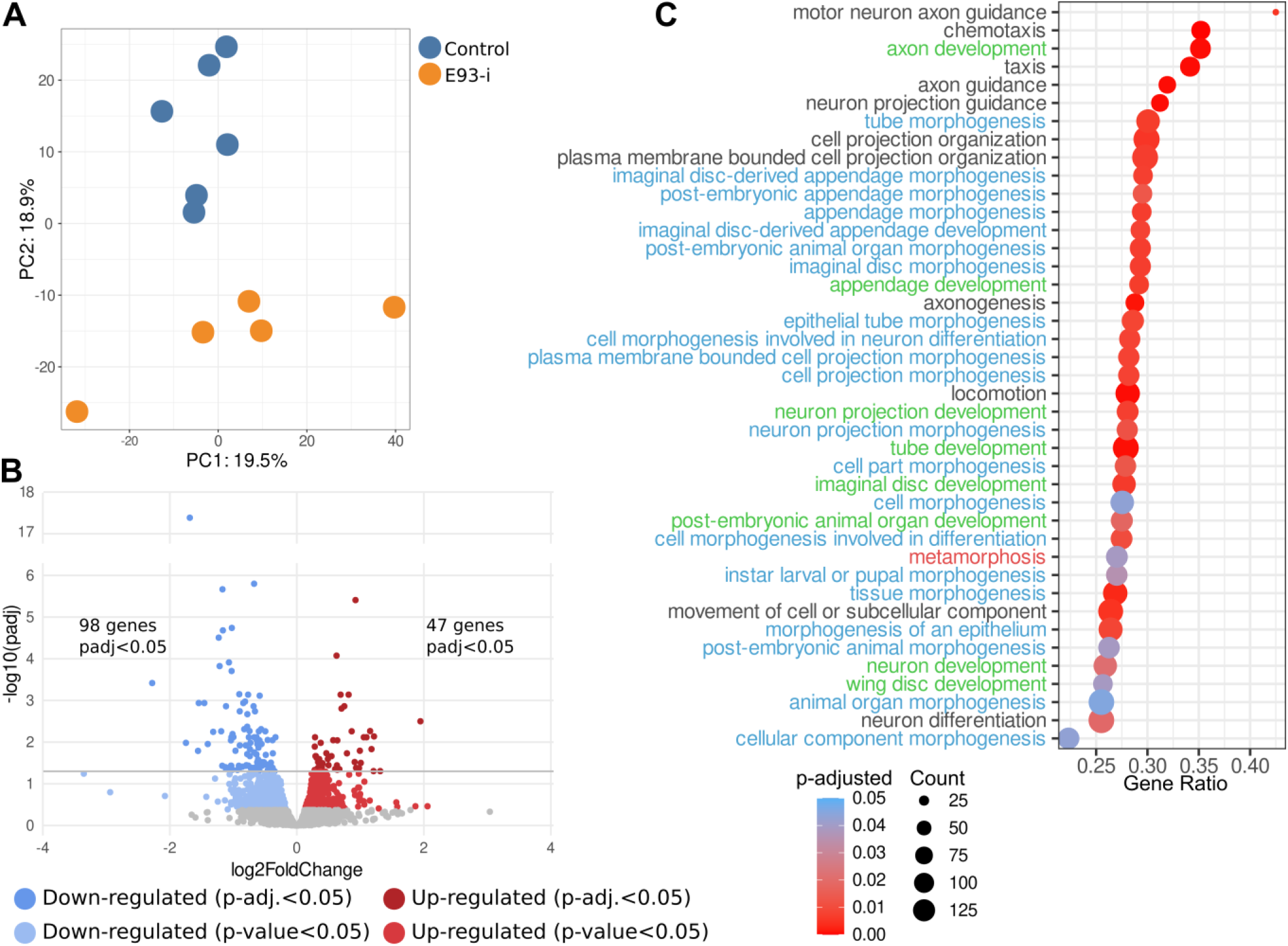
Transcriptomic changes resulting from the depletion of E93 in the embryo of *Blattella germanica*. (*A*) Principal components analysis of the 11 RNA-seq libraries after batch correction (six from controls and five from E93-depleted embryos). (*B*) Volcano plot showing the differentially expressed genes in control and E93-depleted embryos. (*C*) Gene set enrichment analysis (GSEA) showing the top significantly enriched GO terms of biological processes down-regulated after depleting E93 in early embryos. GO terms containing the word “morphogenesis” are in blue, with “development” in green, and “metamorphosis” in red.

These studies revealed that 715 genes tended to be down-regulated, whereas 788 tended to be up-regulated, in E93-depleted samples compared to controls (p-value <0.05). After p-value adjustments for multiple testing, 98 down-regulated and 47 up-regulated genes were found to be statistically significant, with an adjusted p-value <0.05 (Fig. 2*B* and SI Appendix, Dataset S2 and Table S3). Gene set enrichment analysis (GSEA) indicated that the genes down-regulated in E93-depleted embryos are significantly enriched for functions in morphogenesis and development (Fig. 2*C* and 2*D*, and SI Appendix, Fig. S2, Tables S4, S5 and S6).

The 98 genes significantly down-regulated in E93-depleted embryos (SI Appendix, Dataset S2) include gap and pair rule zygotic genes like *Krüppel, even-skipped, fushi tarazu, caudal* and *runt*, which present an expression peak on ED2–ED3 in controls (11). Genes involved in the biosynthesis of ecdysteroids, such as *neverland* and *shade*, which must be involved in the production of the pulse of ecdysone synthesis associated with secretion of the first embryonic cuticle on ED3 (10), also show an expression peak on ED2–ED3, and are significantly down-regulated by E93 depletion. Other genes significantly down-regulated, such as *thisbe, Partner of paired, spalt major, uniflatable*, and *knirps-like*, show an expression pattern starting in ED2 in controls, subsequently maintaining more or less stable levels until the end of embryogenesis. In controls, the 47 genes significantly up-regulated in E93-depleted embryos (SI Appendix, Dataset S2) generally have an expression pattern that shows a sudden decrease in ED2 (11). A number of these may be affected by maternal transcript-clearance mechanisms in the maternal to zygotic transition (MZT), although others might be directly down-regulated by E93. Most of these genes have no orthologues in other species. The few annotated genes include *seven in absentia*, which encodes an E3 ubiquitin ligase, and *starvin*, which encodes a BAG family factor that binds heat shock proteins. In order to validate the results of the differential expression analysis, we measured the expression of nine genes down-regulated and six genes up-regulated in E93-depleted embryos by qRT-PCR. In both groups, the effect of E93 was the same as that observed with RNA-seq (Fig. 3).

**Fig. 3.**
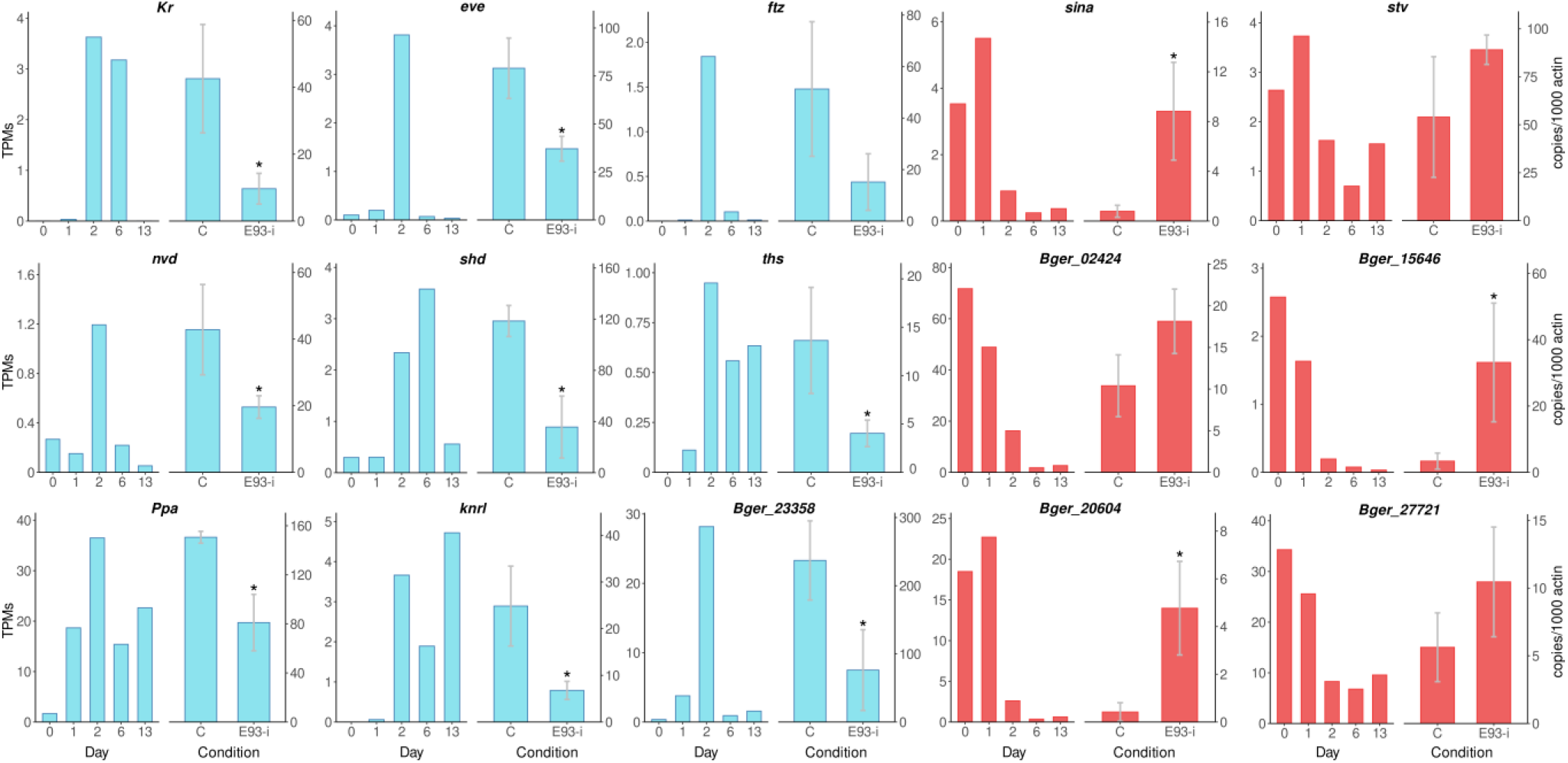
Expression of a selection of genes in E93-depleted (E93-i) embryos of *Blattella germanica*. The following nine genes were selected among the 98 found to be significantly down-regulated (adjusted p-value <0.05) in E93-i samples according to differential expression analysis of transcriptomic data: *Krüppel* (*Kr*), *even-skipped* (*eve*), *fushi tarazu* (*ftz*), *neverland* (*nvd*), *shade* (*shd*), *thisbe* (*ths*) and *Partner of paired* (*Ppa*), *knirps-like* (*knrl*), *Bger_23358*. The following five genes were selected among the 47 found to be significantly up-regulated (adjusted p-value <0.05) according to the same analysis: *seven in absentia* (*sina*), *starvin* (*stv*), *Bger_02424, Bger_15646, Bger_20604*, and *Bger_27721*. The expression profile in 0-, 1-, 2-, 6- and 13-old-day embryogenesis, according to the transcriptomes reported previously (11) (left panel), and the qRT-PCR expression measurements (right panel) in controls (dsMock-treated) and E93-i, are shown for each gene. Each qRT-PCR expression measurement represents 3-5 biological replicates, and results are expressed as copies of the examined transcript per 1000 copies of BgActin-5c mRNA; data are represented as the mean ± SEM; the asterisk indicates statistically significant differences with respect to controls (p-value <0.05), calculated on the basis of the REST (28).

The results suggest that E93 promotes the activation or repression of genes involved in the progression of embryogenesis. We conjecture that the mechanisms underlying this function would be the same that operate in triggering metamorphosis, where E93 acts as a regulator of chromatin accessibility (14, 15). As a result, E93 both opens and closes chromatin in cis regulatory modules during the pre-metamorphic stage, which allows the binding of transcription factors that determine adult morphogenesis (14, 15). We propose that in the early embryo, E93 would contribute to put into motion the genetic program that leads to the development of an adultiform nymph.

### Embryonic expression of *E93* is high in hemimetabolans and low in holometabolans

The high expression of *E93* in the embryo of *B. germanica* contrasts with its very low expression in the embryo of *D. melanogaster* (11). To explore whether this difference is a general trend in hemimetabolan and holometabolan insects, we gathered *E93* expression data for the species where this information was available (SI Appendix, Table S7). To determine whether a given expression in the embryo was “high” or “low”, we compared the embryonic expression with the expression measured in the preadult stage in the same species, in the pupa of holometabolans or in the last nymphal instar in hemimetabolans, that is, the expression associated to adult morphogenesis. For each species, we calculated the log2 ratio between the expression levels of *E93* in the pre-adult stage and the embryo (PA/E ratio). Thus, the higher the PA/E ratio, the lower levels of *E93* expression in the embryo compared to the pre-adult stage. Our results (Fig. 4 and SI Appendix, Table S8) show that the 20 species of holometabolans studied (including Coleoptera, Lepidoptera, Hymenoptera and Diptera) tended to have high PA/E ratios (76.04 ± 40.39). Conversely, the eight typically hemimetabolan studied species (of Ephemeroptera, Blattodea, Orthoptera, Hemiptera Heteroptera and Hemiptera Aphidoidea), have low PA/E ratios (4.43 ± 2.82). Intriguingly, Thysanoptera (*Frankliniella occidentalis* and *Haplothrips brevitubus*) and Hemiptera Coccoidea (*Planococcus kraunhiae*), Aleyrodoidea (*Bemisia tabaci*) and Psylloidea (*Diaphorina citri*) have high PA/E ratios (1928.79 ± 1892.87). These species are currently considered hemimetabolans (3), although some modern authors refer to them as “neometabolans” (17), a name coined by Berlese (18) to categorize the metamorphosis of Coccoidea. We also studied two ametabolan species, namely the Zygentoma *Thermobia domestica*, and the Collembola Entomobryomorpha *Folsomia candida*, which have very low PA/E ratios (0.41 ± 0.11) (Fig. 4). The PA/E ratios for the ametabolans are significantly different to those for holometabolans (p-value = 0.0079) but not different from hemimetabolans (p-value = 0.1455), whereas the ratios for hemimetabolans with respect to neometabolans and holometabolans are also significantly different (p-value = 0.0039 and 0.0009, respectively). The whole data set suggests that high embryonic levels of *E93* expression are generalized in typically hemimetabolan species. According to our observations in *B. germanica*, we conjecture that, in hemimetabolans, E93 plays a crucial role in the activation of the zygotic genome during the MZT, and in the development of the embryo that leads to an adultiform nymph. This contrasts with the embryo of holometabolans, in which *E93* expression is very low and embryogenesis leads to a larval morphology.

**Fig. 4.**
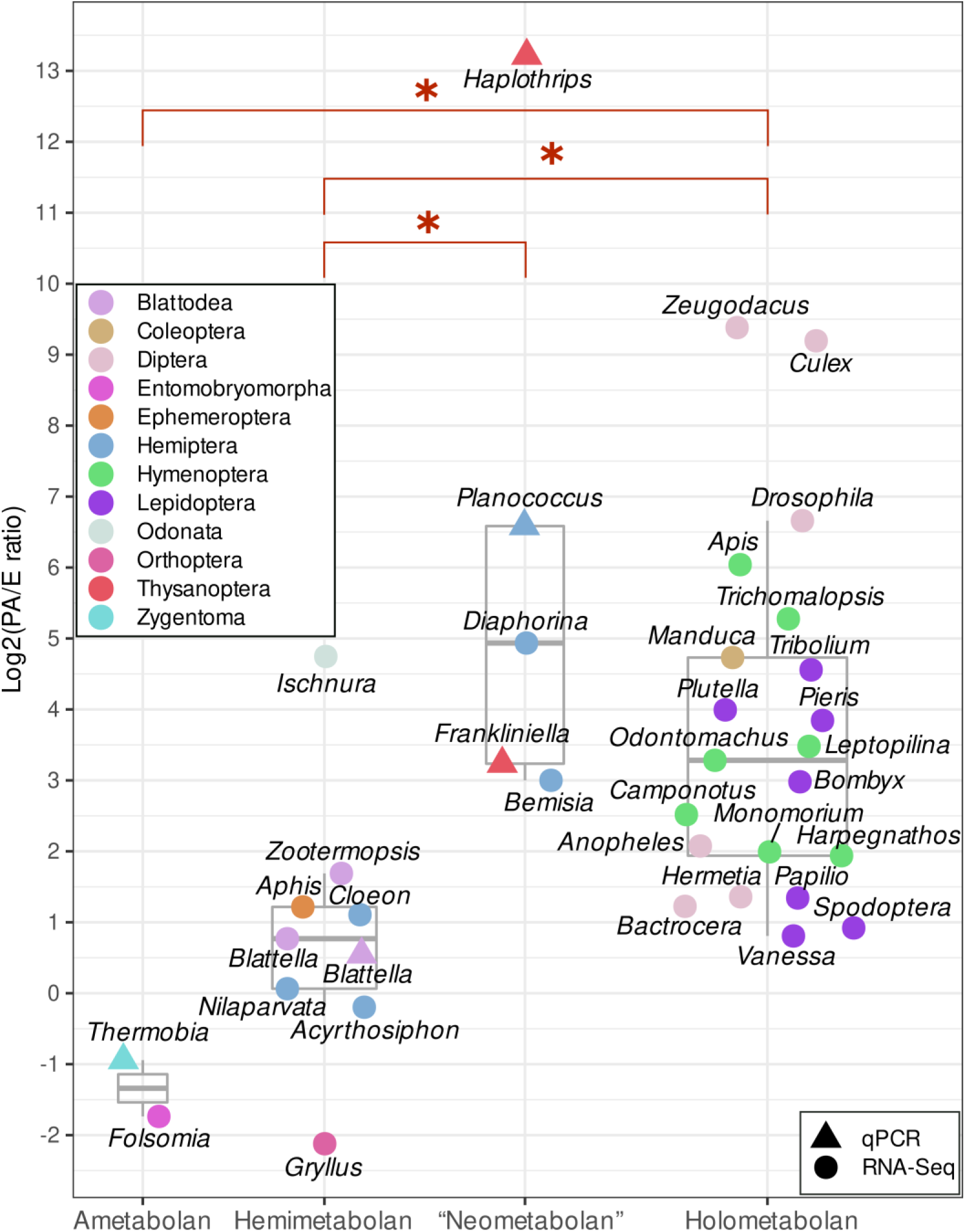
Ratio in log2 scale between the *E93* expression levels in the pre-adult stage and the embryo (PA/E ratio) in different species of ametabolan (*Folsomia candida, Thermobia domestica*), hemimetabolan (*Aphis gossypii, Acyrthosiphon pisum, Blattella germanica, Cloeon dipterum, Gryllus bimaculatus, Ischnura senegalensis, Nilaparvata lugens, Zootermopsis nevadensis*), “neometabolan” (*Bemisia tabaci, Diaphorina citri, Frankliniella occidentalis, Haplothrips brevitubus, Planococcus kraunhiae*), and holometabolan (*Apis mellifera, Anopheles stephensi, Bactrocera dorsalis, Bombyx mori, Camponotus floridanus, Culex quinquefasciatus, Drosophila melanogaster, Harpegnathos saltator, Hermetia illucens, Leptopilina heterotoma, Manduca sexta, Monomorium pharaonis, Odontomachus brunneus, Papilio polytes, Pieris rapae, Plutella xylostella, Spodoptera exigua, Tribolium castaneum, Trichomalopsis sarcophagae, Vanessa tameamea, Zeugodacus cucurbitae*) species. Data are from publicly available RNA-seq data (dots), qRT-PCR measurements (triangles), and colored by the order to which the species belongs. Ratios were calculated as a log2 of the quotient of TPM means for the highest expression sample in pre-adult (nymph or pupa) to embryo stages for each species analyzed. Boxplots indicate the median. Red letters (a, b) indicate statistically (p≤0.05) different groups.

The low embryonic expression of *E93* in Thysanoptera, and in Hemiptera Coccoidea, Aleyrodoidea and Psylloidea might be associated with the unique postembryonic development of these groups, as reviewed elsewhere (3). Thus, that of Thysanoptera includes two or three quiescent stages prior to the adult, in which external features develop and there is considerable internal remodeling. Similarly, Coccoidea show extreme sexual dimorphism, with males passing through two mobile nymphal instars before developing into one or two quiescent instars, then moulting into a winged adult, whereas females undergo a neotenic development. In Psylloidea, the first nymphal instar is mobile but the insect becomes sessile in the second instar, whereas with the imaginal molt, the hind legs dramatically transform into jumping legs. With regard to Aleyrodoidea, the first nymphal instar is mobile but the following nymphal instars are sessile and are shaped as a scale-like, flattened ellipse. The imaginal molt involves extensive tissue destruction and regeneration. The superfamilies Coccoidea, Psylloidea and Aleyrodoidea belong to the Sternorrhyncha, a suborder of Hemiptera that additionally includes the superfamily Aphidoidea, the embryos of which exhibit high *E93* expression levels (Fig. 4) and the post-embryonic development of which does not include quiescent stages. The phylogenetic relationships of insects, in general (19, 20), and of the hemipteran groups with thysanopterans (21, 22) (SI Appendix, Fig. S3), suggest that high *E93* expression levels in the embryo could be an ancestral condition, which might have been reduced in Coccoidea, Psylloidea, Aleyrodoidea and Thysanoptera, as well as in Endopterygota (Holometabola).

Another intriguing result is the low levels of *E93* expression in the embryo of the Odonata, *Ischnura senegalensis*. This may be associated with the morphology of dragonfly nymphs, which is significantly different from that of the adult, including specific and complex nymphal structures, like the protractile labium, that disappear during metamorphosis.

### *E93* is highly expressed in early embryogenesis of the ametabolan *Thermobia domestica*

The data for ametabolan species encouraged us to study the expression pattern of *E93* throughout the life cycle, supposing that ametaboly is the ancestral condition from which hemimetaboly emerged (3). Using the zygentoman *T. domestica*, we observed that *E93* is expressed at very high levels in very early embryo (8% development), decreasing sharply thereafter. During the nymphal period, *E93* expression shows medium values, whereas in the adult female the levels are similar to those of the nymph (Fig. 5*A*). The expression of *E93* in nymphs suggests that it sustains the development of a body plan that has the adult morphology from hatching.

**Fig. 5.**
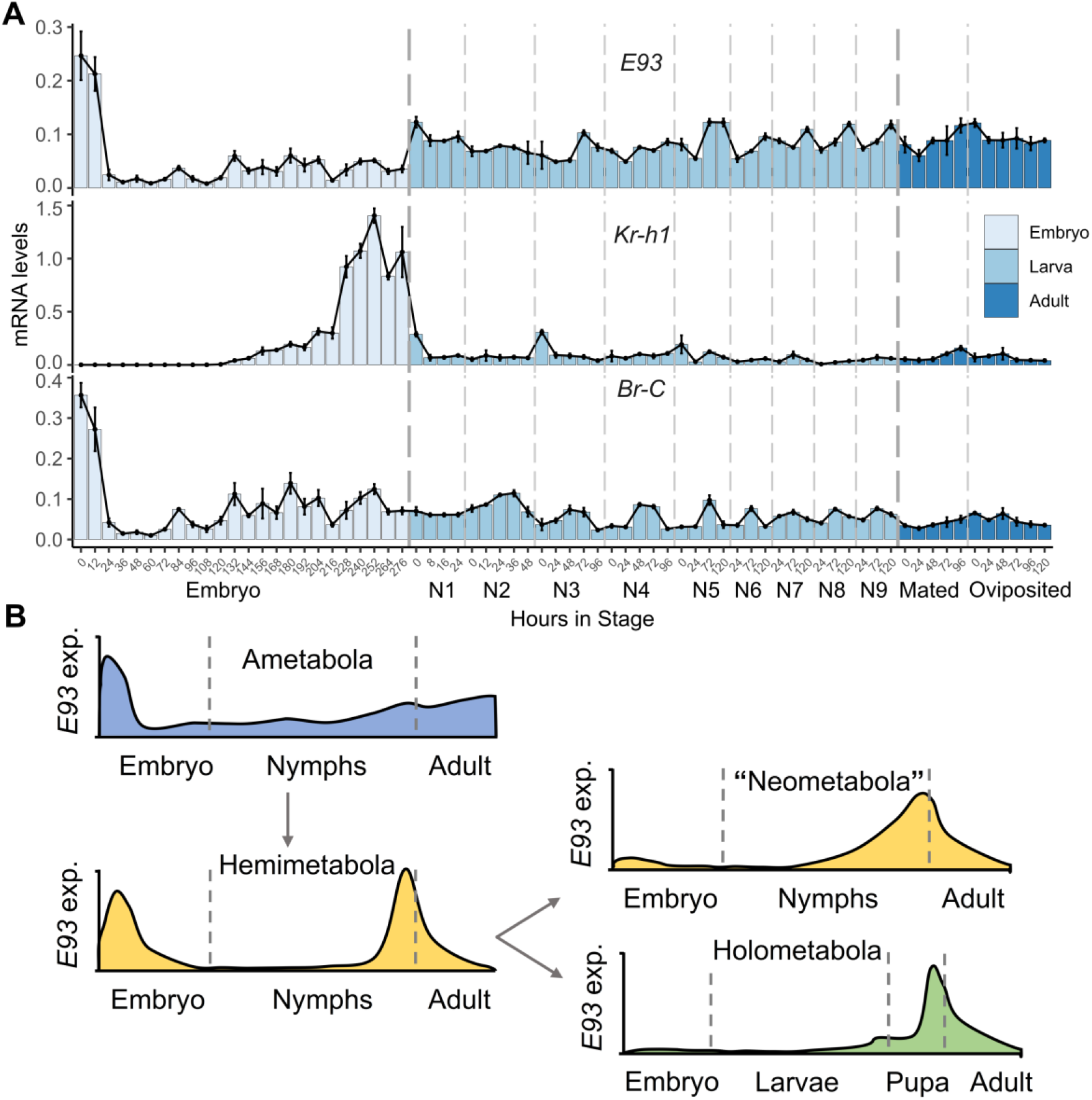
Expression of *E93* in *Thermobia domestica* and hypothesis to explain the evolution of different types of insect metamorphosis in relation to embryonic E93. (*A*). *E93* expression pattern in embryos, nymphs and adults of *T. domestica*, compared with *Kr-h1* and *Br-C* expression; expression was measured by qRT-PCR, and each measurement represents three biological replicates, except for 9th instar nymphs (12 biological replicates); in the case of *Br-C*, we used primers designed in the core region, thus measuring the set of all possible isoforms; mRNA levels were normalized to those of *rp49* and data are represented as the mean ± SEM. (*B*) The evolution of different postembryonic morphologies and types of insect metamorphosis can be explained on the basis of the levels of embryonic *E93* expression. The decrease of *E93* expression in the embryo may have unconstrained the development of the embryo and allowed the formation of new juvenile body forms, and the emergence of new types of metamorphose and life cycles.

We also determined the expression pattern of *Krüppel homolog 1* (*Kr-h1*) to determine any relationship with *E93* expression. In postembryonic development of metamorphosing insects, Kr-h1 is a JH-dependent factor that prevents metamorphosis by repressing *E93* expression, following the MEKRE93 pathway (23, 24). *Kr-h1* expression is very high in the late embryo of *T. domestica* (82–100% development) (Fig. 5*A*), which would correspond to a period of JH production, subsequently falling to very low levels at hatching, with these low levels being maintained throughout the nymphal period. A comparison of the *Kr-h1* expression profile with that for *E93* in the nymphal period does not offer any clear correlations, thus suggesting that there is no reciprocally repressive action between these two factors as occurs in metamorphosing insects (3). Finally, we studied the expression of *Br-C*, obtaining a profile (Fig. 5*A*) strikingly similar to that of *E93* both during embryogenesis and in the nymphal and adult stages. This similarity suggests that Br-C must also be very important at the beginning of embryogenesis in *T. domestica*. In *B. germanica*, maternal RNAi experiments (10, 25) indicate that Br-C has functions in the formation of the germinal band, as well as in other stages of embryo development. Br-C has also been shown to be important in the embryogenesis of another hemimetabolan species, namely the bug *Oncopeltus fasciatus* (9). In contrast, the effects of removing Br-C induced no apparent effects on embryo development of the holometabolan *Bombyx mori* (8).

### Embryonic expression of *E93* and evolution of insect metamorphosis

Our observations suggest that, in direct developing insects (ametabolans and typical hemimetabolans), high levels of embryonic E93 promote activation of the zygotic genome, thereby triggering a type of embryogenesis that culminates in the formation of an adultiform nymph. In holometabolans, low *E93* expression levels in the embryo are associated with a type of embryogenesis that results in the formation of a larva, which is clearly different from the adult. These trends suggest that a reduction of E93 in the embryo may have been instrumental in the emergence of hemimetaboly. We conjecture that a significant decrease of *E93* expression in the embryo would have “unconstrained” the nymphal genetic program, thus allowing alternative developmental programs.

Reduction of *E93* expression in the embryo of Thysanoptera and Hemiptera Sternorrhyncha (except Aphidoidea) could have allowed the morphogenesis of specialized juvenile stages. This would be an independent-convergent innovation in these insects, in the context of their respective processes of adaptation to specific requirements of juveniles. Similarly, a decrease of embryonic *E93* expression could have paved the way for the development of a nymph notably different to the adult in dragonflies. The nymphs of Odonata are secondarily aquatic, thus the above process may have been associated with the colonization of freshwater habitats by juveniles, including specific adaptations to predatory habits.

Taken together, our data suggest that the last common ancestor of insects, which emerged in the early Paleozoic, would have high levels of *E93* expression in the embryo thus resulting in the development of nymphs very similar to the adult, whereas during the entire postembryonic life, *E93* expression would remain at medium levels. In the early-middle Devonian, hemimetaboly emerged, which is associated with the emergence of wings and the innovation of the final molt (3). The suppression of *E93* expression during the nymphal period and its concentration in the preadult stage appears to be consubstantial with the innovation of the final moult associated to hemimetaboly (26). Later, during the early Carboniferous, holometaboly emerged, involving the formation of new kinds of juvenile stages: the larva and the pupa. The attenuation of *E93* expression in the embryo could have been a key step in allowing the launch of the larval genetic program, in the context of the new type of metamorphosis. Significant and independent decreases of *E93* expression in the embryo of Odonata, Thysanoptera, and Hemiptera Coccoidea, Psylloidea and Aleyrodoidea, would have allowed the development of modified juvenile stages adapted to specific ecophysiological conditions (Fig. 5*B*).

As noted above, in the study of insect metamorphosis, especially as regards the regulatory and evolutionary aspects, embryo development has been largely overlooked. However, it is obvious that the type of embryogenesis determines the morphology of the juveniles and the type of metamorphosis. Thus, the study of the evolution of the different types of insect embryogenesis is key to understanding the evolution of the different types of insect metamorphosis and life cycles. There is nothing new under the sun. *Ex ovo omnia*, as Harvey said in the XVII Century.

## Materials and methods

A detailed description is given in SI Appendix. In brief, the rearing methods of *B. germanica* and *T. domestica* were as reported previously (25, 27). Methods for RNA extraction and reverse transcription to complementary DNA and qRT-PCR were essentially as described elsewhere (25). Maternal RNAi was performed as reported earlier (25). Embryo morphology was studied with a Zeiss DiscoveryV8 stereomicroscope, and a Carl Zeiss-AXIO IMAGER.Z1 bright-field microscope. Preparation and sequencing of mRNA libraries, and their bioinformatic study was performed as described previously (11). To estimate the expression levels of *E93* across hexapods, publicly available RT-qPCR and RNA-seq datasets were used.

## Data Availability

All the RNA-seq datasets of *B. germanica* generated are publicly available at NCBI-SRA under the BioProject number PRJNA865117.

## ACKNOWLEDGEMENTS

The work was supported by the Spanish Agencia Estatal de Investigación (grant number PID2019-104483GB-I00 / AEI / 10.13039/501100011033; CGL2015-64727-P; and CGL2012-36251, to X.B), the Catalan Government (2017 SGR 1030, to X.B), and the European Fund for Economic and Regional Development (FEDER funds, to X.B), as well as the Ministry of Education, Culture, Sports, Science and Technology of Japan (Grants-in-Aid for Scientific Research, KAKENHI, numbers. 20H02999 and 17H03943, to T.D.) and JSPS Bilateral Joint Research Projects (JPJSBP120209917, to T.D.). Part of this research has been financed (to G.Y.) by the Faculty of Biochemistry, Biophysics and Biotechnology at Jagiellonian University (Poland), under the Strategic Programme Excellence Initiative Thanks are also due to Maria-Dolors Piulachs and Jose Luis Maestro for helpful discussions.

