## Supplemental information for "Reduction of embryonic *E93* expression as a key factor for the evolution of insect metamorphosis"

### Supporting information

Supplementary Materials and methods

Table S1. Libraries prepared to study the influence of E93 on other genes.

Table S2. Reads mapping to the reference genome.

Table S3. Differentially expressed genes. Summary of the results.

Table S4. Significantly enriched pathways (Kegg analysis)

Table S5.  Significantly enriched pathways (GO terms).

Table S6. Enrichment analysis. Summary of the results.

Table S7. Species where E93 expression in embryos and pre-adult stage was obtained with publicly available data.

Table S8. E93 Ratios in the analyzed species.

Table S9. Primer sequences used for qRT-PCR and RNAi experiments.

Figure S1. Principal Component Analysis of all available libraries.

Figure S2. Gene Set Enrichment Analysis (GSEA) of the RNA-seq from the E93 depletion experiment.

Figure S3. Phylogeny the Hemiptera Sternorrhyncha, and relationships with Thysanoptera and Psocodea.

### Materials and Methods

**Insect rearing.** *B. germanica* specimens used in the experiments were obtained from a colony reared in the dark at 30 ±1ºC and 60-70% r.h. The experimental insects were carbon dioxide-anaesthetized prior to dissections and tissue sampling. *T. domestica* specimens were from a colony reared at 37 ±1ºC and 60-80% r.h. as described previously [1]. To stage the embryos, the cotton in a rearing container (on which eggs were laid), was replaced every four hours to obtain newly laid eggs. To stage the nymphs, they were individually reared in a plastic container, marked with a felt-tip pen on the dorsal side, and the occurrence of molt was monitored every 4 h between the first and the fifth instar, and every 12 h between sixth to ninth instar, in order to obtain newly molted nymphs. Finally, newly molted female adults (within 12 h of adult molt) were individually placed in a plastic container with six male adults for 12 h, then only females to which spermatophores were attached during this period were used for the experiments.

**RNA Extraction and  reverse transcription to cDNA.** In the case of *B. germanica*, RNA extractions were carried out with the Gen Elute Mammalian Total RNA kit (Sigma-Aldrich). Then RNA extracts were treated with DNase (Promega) and reverse transcribed with Superscript II reverse transcriptase (Invitrogen) and random hexamers (Promega). In *T. domestica*, total RNA was extracted using an RNeasy Plus Mini kit (Qiagen), and was used to synthesize cDNA with a ReverTra Ace qPCR RT Master Mix with gDNA Remover kit (ToYoBo).

**Determination of mRNA levels with quantitative real-time PCR.** In the case of *B. germanica*, quantitative real time PCR (qRT-PCR) reactions were carried out in triplicate in an iQ5 Real-Time PCR Detection System (Bio-Rad Laboratories), using SYBR®Green (Power SYBR® Green PCR Master Mix; Applied Biosystems). The gene *actin-5c* was used as a reference. For T. *domestica*, we used a Thermal Cycler Dice Real Time System (TaKaRa), using THUNDERBIRD SYBR qPCR Mix (ToYoBo). The gene *rp49* served as the internal reference. A control without template was included in all batches. The primers used for each transcript measured are detailed in Table S9 The efficiency of each primer set was first validated by constructing a standard curve through four serial dilutions. In *B. germanica*, for comparison of controls and E93-depleted samples, we followed a method based in Ct (threshold-cycle) according to the Pfaffl mathematical model [2], simplifying to $2^{\Delta\Delta Ct}$ because the calculated efficiency values for studied genes and actin-5c amplicons were always within the range of 95 to 100%; therefore, no correction for efficiency was used in further calculations. Results are given as copies of mRNA per 1,000 copies of actin-5c mRNA. Statistical differences between groups were tested by the REST 2008 program (Relative Expression Software Tool V 2.0.7; Corbett Research) [2]. In *T. domestica*, expression levels were quantified via an absolute quantification method using a dilution series of a standard plasmid with an insert of a fragment of the same DNA that is being quantified in samples. Results are given as relative amounts of *rp49*.

**Maternal RNA interference.** Detailed procedures to prepare the dsRNA for RNAi were as described previously [3], and the corresponding E93 primers are described in Table S9. A dsRNA from *Autographa californica* nucleopoydrovirus was used for control treatments (dsMock). Maternal RNAi treatments were carried out essentially as previously reported [4]. A volume of 1 µl of dsRNA solution (3 µg/µl) was injected into the abdomen of 5-dayold adult females. Then, the effects of the treatment were examined in the embryos of the first ootheca on selected days after their formation.

**Examination of embryos.** To examine the embryos microscopically, the oothecae were opened after 5 min in a water bath at 95°C and the embryos were dechorionated and individualized. Then, they were fixed in 4% paraformaldehyde, permeabilized in PBS+0.2% Tween (PBT) and incubated for 10 min in 1 µg/ml DAPI in PBT. They were then mounted in Mowiol 4-88 (Calbiochem) and examined and photographed using epifluorescence with an AxioImager Z1 microscope (ApoTome System, Zeiss). Embryo stages were established using the criteria and nomenclature of Tanaka [5].

**Preparation and sequencing of mRNA libraries.** We prepared and sequenced 6 mRNA libraries of a pool of 2-day-old oothecae produced by mated females that had been treated with dsE93 on adult day 5. Six equivalent control libraries (females treated with dsMock) were also prepared and sequenced. Total RNA was extracted using the GenElute Mammalian Total RNA kit (Sigma) following the manufacturer's protocol. Up to 10 µg of total RNA from pooled samples were used to prepare the libraries. mRNA was obtained from RNA samples using NEBNext® Poly(A) mRNA Magnetic Isolation Module (NEB; Herts, UK). After library preparation (NEBNext® Ultra™ II RNA

Library Prep Kit for Illumina, NEB), cDNA was sequenced in PE150 (paired ends mode, 150 bp) using NextSeq 500 (Illumina; San Diego, CA, USA) next-generation sequencing platform. In total 12 RNA libraries were sequenced in two batches (6 samples per batch).

**Reads trimming, quality control, and reads mapping.** The raw sequences were subjected to adaptor removal and quality control with Trimmomatic (v.0.39) [6] using TRueSeq3-PE.fa.2:30:10.8 adapters and default parameters. Reads quality was determined with the FastQC tool (v.0.11.9) [7]. Reads were then mapped to the *Blattella germanica* genome using STAR (v.2.7.9a) [8] with the default parameters. MultiQC (v.1.11) [9] was used to summarize mapping and reads quality data.

**Reads count, batch effect removal, differential expression, and gene set enrichment analysis.** Reads mapped to genes were counted in R (v.4.1.2) using the featureCounts algorithm implemented in the Rsubread package (v. 2.8.1 ) [10]. To remove the batch effect sva (v.3.42.0) [11] and limma (v.3.50.0) [12] R packages were used. Differential expression analysis (DEA) and gene set enrichment analysis (GSEA) were conducted using DESeq2 (v.1.34.0) [13] and clusterProfiler (v.4.2.1) [14] R packages, respectively. Scripts used in the data analysis are publicly available at: https://github.com/ylla-lab/Embryonic_E93.

**Sampling *E93* expression data in selected hexapods.** To estimate the expression levels of *E93* across hexapods, publicly available RT-qPCR and RNA-seq datasets were used. RT-qPCR data were obtained from two species of Thysanoptera (*Frnakliniella occidentalis* and *Haplothrips brevitubus*) [15], a species of Hemiptera Coccoidea (*Planococcus krauhniae*) [16], and a species of Odonata (*Ischnura senegalensis*) [17] (Table S7). The RNA-seq data was obtained from NCBI as follows. The NCBI's SRA database was queried for RNA-seq datasets of hexapods (taxon id 6960) generated with Illumina (with the exceptions of *Folsomia candida* and *Aphis gossypii* for which BGI-seq was also used), in fastq format and excluding "mirna-seq". The terms "embryo" and "egg" were used to query embryonic RNA-seq data, and "nymph", "juvenile", and "pupa", for the juvenile stages. Metadata for the 9,925 RNA-seq datasets of embryo, 16,203 of nymph, and 1,058 of pupa, were downloaded. Species for which no annotated genome was available were discarded. To the list of 263 Hexapod species with the available genome in NCBI as of January 2022, *Gryllus bimaculatus* was added given that the annotated genome was available elsewhere [18]. The species with the annotated genome were looked at in the downloaded RNA-seq metadata. Those species with annotated genome that had RNA-seq data from embryo, and pupa or nymph, were pre-selected. A manual review of the pre-selected datasets was performed to remove non-mRNA libraries (e.g. RIP-seq), inappropriate samples source (e.g. cell cultures), samples from treated or mutant specimens, and unrepresentative tissues when multiple tissues were available. For hemiemtabolans, the last available nymphal instar was selected, and validated that it was the pre-adult nymph (out of 11 species, 2 of them, *Zootermopsis nevadensis* and *Bemisia tabaci*, could not be confirmed that the last available juvenile RNA-seq sample was indeed the pre-adult due to lack of metadata information). Manual filtering resulted in a list of 848 RNA-seq datasets from 33 Hexapoda species (Table S7) that were downloaded and decompressed with prefetch and fastq-dump utils from SRA-Toolkit (https://trace.ncbi.nlm.nih.gov/Traces/sra/sra.cgi?view=software). The RNA-seq reads were subjected to adaptor removal with TrimGalore (v.0.6.7) (https://github.com/FelixKrueger/TrimGalore) with default parameters. Then, reads were mapped to the corresponding genomes with STAR. Genomes and annotations were downloaded from NCBI, except those of *G. bimaculatus* and *B. germanica*, for which we used the original files from publication [18,19]. *E93* gene in each species was identified using the *Drosophila melanogaster* E93 transcript sequence (Eip93F; NM_001104395.2) as a query against each species protein database using BLASTX. For *Trichomalopsis sarcophagae* and *Megalopta genalis* no significant hits were found. Using TBLASTX against the genome assembly, an *E93* homolog was identified and annotated in the *T. sarcophagae* genome (scaffold NNAY01002743.1:14775-19230), but no significant matches were found for *M. genalis* which had to be excluded from further analysis. The featureCounts algorithm implemented in the Rsubread package [10] was used to generate the table of counts for each species, which were then normalized as Transcript per Million (TPMs). For each species, the pre-adult to embryo ratio (PA/E ratio) was calculated as follows. The RNA-seq dataset with the highest E93 TPMs for embryo and pre-adult were selected, if the selected dataset had biological replicates available those were averaged. Then, the TPMs of pre-adult were divided by the TPMs of the embryo and log2-transformed for the data visualizations.

**Table S1. Libraries prepared to study the influence of E93 on other genes.** Early embryo E93 mRNA was depleted through maternal RNAi, and RNA was sampled on day 2 of embryo development

| Library | Index | Treatment |
|---|---|---|
| 1 | 16 | dsMock (control) |
| 2 | 18 | dsMock (control) |
| 3 | 20 | dsMock (control) |
| 6 | 23 | dsE93 |
| 8 | 25 | dsE93 |
| 9 | 27 | dsE93 |
| C9 | 15 | dsMock (control) |
| C11 | 19 | dsMock (control) |
| C12 | 27 | dsMock (control) |
| T7 | 5 | dsE93 |
| T6 | 3 | dsE93 |
| T14 | 9 | dsE93 |

**Table S2. Reads mapping to the reference genome.**

| Library | Treatment | Total reads | Uniquely mapped | Uniquely mapped percent | Multi mapped | Multi mapped percent | Multi mapped toomany | Multi mapped toomany percent | Unmapped tooshort percent | Unmapped other percent | Unmapped tooshort | Unmapped other |
|---|---|---|---|---|---|---|---|---|---|---|---|---|
| 1 | dsMock (control) | 26710703 | 24313538 | 91.03 | 1055496 | 34759.00 | 33799 | 0.13 | 27851.00 | 0.14 | 1270502.00 | 37368.00 |
| 2 | dsMock (control) | 29885886 | 27185636 | 90.96 | 1188340 | 35855.00 | 35802 | 0.12 | 29677.00 | 0.13 | 1437263.00 | 38845.00 |
| 3 | dsMock (control) | 27486910 | 24920143 | 90.66 | 1105500 | 44596.00 | 29047 | 0.11 | 44870.00 | 0.1 | 1404730.00 | 27490.00 |
| 6 | dsE93 | 26435547 | 24123313 | 91.25 | 1065345 | 44624.00 | 31895 | 0.12 | 17989.00 | 0.11 | 1185940.00 | 29054.00 |
| 8 | dsE93 | 28133923 | 25736622 | 91.48 | 1129862 | 44596.00 | 29627 | 0.11 | 44624.00 | 0.1 | 1209680.00 | 28132.00 |
| 9 | dsE93 | 27798417 | 25079532 | 90.22 | 1113942 | 44565.00 | 28141 | 0.1 | 21306.00 | 0.09 | 1551773.00 | 25029.00 |
| C11 | dsMock (control) | 32117037 | 28927803 | 90.07 | 1379147 | 47209.00 | 49591 | 0.15 | 13271.00 | 0.13 | 1718808.00 | 41688.00 |
| C12 | dsMock (control) | 29047807 | 25679314 | 88.4 | 1266886 | 13241.00 | 44582 | 0.15 | 35582.00 | 0.11 | 2025066.00 | 31959.00 |
| C9 | dsMock (control) | 29664738 | 26533447 | 89.44 | 1263026 | 46113.00 | 44906 | 0.15 | 44598.00 | 0.13 | 1784816.00 | 38543.00 |
| T14 | dsE93 | 21303180 | 19103816 | 89.68 | 915618 | 44624.00 | 40180 | 0.19 | 44747.00 | 0.13 | 1215836.00 | 27730.00 |
| T6 | dsE93 | 31822342 | 25116900 | 78.93 | 1171151 | 24898.00 | 37631 | 0.12 | 17.17 | 0.1 | 5464832.00 | 31828.00 |

**Table S3. Differentially expressed genes**. Summary of the results.

| **Statistical criteria** | **Total** | **Up-regulated** | **Down-regulated** |
|---|---|---|---|
| padj <0.05 | 145 | 47 | 98 |
| padj <0.05 + \|log2Fvh\|>1 | 36 | 9 | 27 |
| p<0.05 | 1503 | 788 | 715 |
| p<0.05 + \|log2Fvh\|>1 | 217 | 64 | 153 |

**Table S4. Significantly enriched pathways** (Kegg analysis)

| ID | Description | Set Size | Enrichment Score | NES | pvalue | p.adjust | qvalues | Rank | Leading edge |
|---|---|---|---|---|---|---|---|---|---|
| dme03010 | Ribosome | 98 | 0.5903152 | 2.497943301 | 1.00E-10 | 1.00E-10 | 1.07E-08 | 1846 | tags=80%. list=35%. signal=53% |
| dme03040 | Spliceosome | 87 | 0.47390276 | 1.950804329 | 2.89E-05 | 2.89E-05 | 0.001241265 | 2486 | tags=78%. list=47%. signal=42% |
| dme00190 | Oxidative phosphorylation | 73 | 0.50577936 | 2.033450033 | 3.47E-05 | 3.47E-05 | 0.001241265 | 1642 | tags=56%. list=31%. signal=39% |
| dme03008 | Ribosome biogenesis in eukaryotes | 51 | 0.5543937 | 2.061946146 | 5.17E-05 | 5.17E-05 | 0.001388459 | 1671 | tags=73%. list=31%. signal=50% |
| dme00830 | Retinol metabolism | 4 | -0.91457454 | -1.67124798 | 0.003225619 | 0.003225619 | 0.063941812 | 186 | tags=100%. list=3%. signal=97% |
| dme00350 | Tyrosine metabolism | 7 | -0.83870592 | -1.791502223 | 0.003824734 | 0.003824734 | 0.063941812 | 31 | tags=57%. list=1%. signal=57% |
| dme01100 | Metabolic pathways | 564 | -0.28306847 | -1.333016568 | 0.004168755 | 0.004168755 | 0.063941812 | 605 | tags=15%. list=11%. signal=15% |
| dme00970 | Aminoacyl-tRNA biosynthesis | 32 | 0.52121008 | 1.745326805 | 0.008536759 | 0.008536759 | 0.107270776 | 2067 | tags=72%. list=39%. signal=44% |
| dme00981 | Insect hormone biosynthesis | 6 | -0.83115852 | -1.700588605 | 0.008991815 | 0.008991815 | 0.107270776 | 280 | tags=50%. list=5%. signal=47% |
| dme00020 | Citrate cycle (TCA cycle) | 21 | 0.57348647 | 1.730560677 | 0.011190317 | 0.011190317 | 0.111640748 | 65 | tags=14%. list=1%. signal=14% |
| dme04512 | ECM-receptor interaction | 10 | -0.72287203 | -1.675358727 | 0.013594603 | 0.013594603 | 0.111640748 | 526 | tags=70%. list=10%. signal=63% |
| dme00330 | Arginine and proline metabolism | 15 | -0.64389125 | -1.661991011 | 0.013760676 | 0.013760676 | 0.111640748 | 456 | tags=40%. list=9%. signal=37% |
| dme00561 | Glycerolipid metabolism | 16 | -0.64915405 | -1.718917026 | 0.014336336 | 0.014336336 | 0.111640748 | 1599 | tags=75%. list=30%. signal=53% |
| dme03013 | Nucleocytoplasmic transport | 71 | 0.38799292 | 1.56446478 | 0.01457555 | 0.01457555 | 0.111640748 | 2359 | tags=63%. list=44%. signal=36% |
| dme00040 | Pentose and glucuronate interconversions | 8 | -0.75755639 | -1.673991255 | 0.015596869 | 0.015596869 | 0.111640748 | 763 | tags=50%. list=14%. signal=43% |
| dme00250 | Alanine. aspartate and glutamate metabolism | 20 | -0.56403988 | -1.589796448 | 0.023195998 | 0.023195998 | 0.155657357 | 527 | tags=35%. list=10%. signal=32% |

| | | | | | | | | | |
|---|---|---|---|---|---|---|---|---|---|
| dme00440 | Phosphonate and phosphinate metabolism | 4 | 0.82607607 | 1.519770456 | 0.026929287 | 0.026929287 | 0.170079708 | 430 | tags=25%. list=8%. signal=23% |
| dme04070 | Phosphatidylinositol signaling system | 41 | -0.4594503 | -1.511502736 | 0.032144851 | 0.032144851 | 0.174628595 | 604 | tags=41%. list=11%. signal=37% |
| dme00053 | Ascorbate and aldarate metabolism | 5 | -0.82224137 | -1.591074898 | 0.035282339 | 0.035282339 | 0.174628595 | 156 | tags=80%. list=3%. signal=78% |
| dme04142 | Lysosome | 45 | -0.44680598 | -1.503755355 | 0.038027296 | 0.038027296 | 0.174628595 | 1125 | tags=40%. list=21%. signal=32% |
| dme04320 | Dorso-ventral axis formation | 20 | -0.53771861 | -1.515607624 | 0.038345261 | 0.038345261 | 0.174628595 | 515 | tags=35%. list=10%. signal=32% |
| dme01230 | Biosynthesis of amino acids | 34 | 0.43120965 | 1.472481 | 0.038630894 | 0.038630894 | 0.174628595 | 170 | tags=18%. list=3%. signal=17% |
| dme03022 | Basal transcription factors | 30 | 0.45545915 | 1.494353925 | 0.038991931 | 0.038991931 | 0.174628595 | 1986 | tags=50%. list=37%. signal=32% |
| dme00230 | Purine metabolism | 45 | -0.44570886 | -1.500062923 | 0.039034627 | 0.039034627 | 0.174628595 | 673 | tags=27%. list=13%. signal=24% |
| dme00562 | Inositol phosphate metabolism | 34 | -0.47420798 | -1.503890004 | 0.041411159 | 0.041411159 | 0.177850032 | 1136 | tags=44%. list=21%. signal=35% |
| dme00592 | Alpha-Linolenic acid metabolism | 3 | -0.86267668 | -1.451895741 | 0.046535808 | 0.046535808 | 0.192172162 | 449 | tags=67%. list=8%. signal=61% |

**Table S5. Significantly enriched pathways** (GO terms)

| Category | ID | Description | Set Size | Enrichment Score | NES | pvalue | p.adjust | qvalues | Rank | Leading edge |
|---|---|---|---|---|---|---|---|---|---|---|
| MF | GO:0003723 | RNA binding | 481 | 0.454223944 | 2.245536199 | 1.00E-10 | 2.17E-08 | 1.92E-08 | 4011 | tags=64%. list=36%. signal=42% |
| MF | GO:0003735 | structural constituent of ribosome | 126 | 0.608614897 | 2.568094441 | 1.00E-10 | 2.17E-08 | 1.92E-08 | 3136 | tags=72%. list=28%. signal=52% |
| MF | GO:0140098 | catalytic activity. acting on RNA | 174 | 0.471729913 | 2.077150125 | 1.76E-08 | 2.55E-06 | 2.26E-06 | 3424 | tags=61%. list=31%. signal=43% |
| MF | GO:0140640 | catalytic activity. acting on a nucleic acid | 268 | 0.395972251 | 1.847256405 | 2.18E-07 | 2.36E-05 | 2.09E-05 | 3424 | tags=49%. list=31%. signal=35% |
| MF | GO:0008168 | methyltransferase activity | 84 | 0.533452795 | 2.09142468 | 6.78E-06 | 0.0005887 | 0.000521162 | 3411 | tags=60%. list=31%. signal=41% |
| MF | GO:0140101 | catalytic activity. acting on a tRNA | 62 | 0.563032879 | 2.08266484 | 1.01E-05 | 0.00073336 | 0.000649229 | 3424 | tags=74%. list=31%. signal=51% |
| MF | GO:0016741 | transferase activity. transferring one-carbon groups | 89 | 0.496701949 | 1.964032588 | 2.36E-05 | 0.00146221 | 0.00129446 | 3411 | tags=57%. list=31%. signal=40% |
| MF | GO:0005198 | structural molecule activity | 198 | 0.384098089 | 1.719952955 | 2.98E-05 | 0.00161448 | 0.001429267 | 3040 | tags=51%. list=28%. signal=38% |
| MF | GO:0005509 | calcium ion binding | 112 | -0.4891187 | -1.832376385 | 7.84E-05 | 0.00378043 | 0.00334673 | 2313 | tags=40%. list=21%. signal=32% |
| MF | GO:0008757 | S-adenosylmethionine-dependent methyltransferase activity | 63 | 0.515285306 | 1.911579734 | 0.000228664 | 0.009924 | 0.008785494 | 3593 | tags=60%. list=33%. signal=41% |
| MF | GO:0008173 | RNA methyltransferase activity | 37 | 0.583086507 | 1.94591058 | 0.000426886 | 0.01684258 | 0.014910363 | 3411 | tags=76%. list=31%. signal=52% |
| MF | GO:0019843 | rRNA binding | 30 | 0.63497581 | 2.026423424 | 0.000500682 | 0.01810798 | 0.016030594 | 2799 | tags=70%. list=25%. signal=52% |
| MF | GO:0008061 | chitin binding | 13 | -0.79212453 | -1.933285372 | 0.000553381 | 0.01847441 | 0.01635498 | 818 | tags=62%. list=7%. signal=57% |
| MF | GO:0070851 | growth factor receptor binding | 11 | -0.80333041 | -1.880977489 | 0.000789317 | 0.02446884 | 0.021661716 | 901 | tags=55%. list=8%. signal=50% |
| MF | GO:0000981 | DNA-binding transcription factor activity. RNA polymerase II-specific | 175 | -0.39843009 | -1.578599908 | 0.001113707 | 0.03222325 | 0.028526524 | 1716 | tags=26%. list=16%. signal=22% |
| MF | GO:0003743 | translation initiation factor activity | 35 | 0.58469446 | 1.933299042 | 0.001258158 | 0.03412752 | 0.030212335 | 3996 | tags=80%. list=36%. signal=51% |
| MF | GO:0050839 | cell adhesion molecule binding | 40 | -0.58762776 | -1.838815375 | 0.001388519 | 0.03544806 | 0.031381379 | 2230 | tags=50%. list=20%. signal=40% |
| BP | GO:0006364 | rRNA processing | 116 | 0.625467614 | 2.606882033 | 1.00E-10 | 2.11E-08 | 1.99E-08 | 3092 | tags=77%. list=28%. signal=56% |
| BP | GO:0006396 | RNA processing | 453 | 0.482958076 | 2.374223125 | 1.00E-10 | 2.11E-08 | 1.99E-08 | 3597 | tags=62%. list=33%. signal=44% |
| BP | GO:0006412 | translation | 324 | 0.464006215 | 2.209785041 | 1.00E-10 | 2.11E-08 | 1.99E-08 | 3547 | tags=60%. list=32%. signal=42% |
| BP | GO:0006518 | peptide metabolic process | 388 | 0.405766966 | 1.970500023 | 1.00E-10 | 2.11E-08 | 1.99E-08 | 3559 | tags=55%. list=32%. signal=39% |
| BP | GO:0016072 | rRNA metabolic process | 124 | 0.61726463 | 2.593716806 | 1.00E-10 | 2.11E-08 | 1.99E-08 | 3092 | tags=75%. list=28%. signal=55% |

| | | | | | | | | | | |
|---|---|---|---|---|---|---|---|---|---|---|
| BP | GO:0022613 | ribonucleoprotein complex biogenesis | 214 | 0.58333525 | 2.644883243 | 1.00E-10 | 2.11E-08 | 1.99E-08 | 3111 | tags=67%. list=28%. signal=49% |
| BP | GO:0034470 | ncRNA processing | 207 | 0.563120476 | 2.548822156 | 1.00E-10 | 2.11E-08 | 1.99E-08 | 3573 | tags=74%. list=32%. signal=51% |
| BP | GO:0034660 | ncRNA metabolic process | 253 | 0.540248957 | 2.498863155 | 1.00E-10 | 2.11E-08 | 1.99E-08 | 3573 | tags=70%. list=32%. signal=48% |
| BP | GO:0042254 | ribosome biogenesis | 161 | 0.617737052 | 2.705160029 | 1.00E-10 | 2.11E-08 | 1.99E-08 | 3092 | tags=74%. list=28%. signal=54% |
| BP | GO:0043043 | peptide biosynthetic process | 351 | 0.439950111 | 2.112641967 | 1.00E-10 | 2.11E-08 | 1.99E-08 | 3853 | tags=63%. list=35%. signal=42% |
| BP | GO:0043604 | amide biosynthetic process | 372 | 0.412065154 | 1.990968771 | 1.00E-10 | 2.11E-08 | 1.99E-08 | 3853 | tags=61%. list=35%. signal=41% |
| BP | GO:0140053 | mitochondrial gene expression | 101 | 0.604420887 | 2.464373516 | 1.13E-10 | 2.19E-08 | 2.07E-08 | 3104 | tags=68%. list=28%. signal=50% |
| BP | GO:0032543 | mitochondrial translation | 81 | 0.642325428 | 2.507370016 | 2.32E-10 | 4.15E-08 | 3.92E-08 | 3042 | tags=72%. list=28%. signal=52% |
| BP | GO:0043603 | cellular amide metabolic process | 429 | 0.369478458 | 1.808591819 | 6.66E-09 | 1.10E-06 | 1.04E-06 | 3559 | tags=53%. list=32%. signal=37% |
| BP | GO:0006397 | mRNA processing | 227 | 0.419501885 | 1.917770289 | 9.99E-08 | 1.55E-05 | 1.46E-05 | 4157 | tags=64%. list=38%. signal=41% |
| BP | GO:0002181 | cytoplasmic translation | 97 | 0.547849187 | 2.214320536 | 1.67E-07 | 2.42E-05 | 2.28E-05 | 3783 | tags=79%. list=34%. signal=53% |
| BP | GO:0006399 | tRNA metabolic process | 103 | 0.527112433 | 2.151482038 | 3.25E-07 | 4.44E-05 | 4.19E-05 | 3573 | tags=71%. list=32%. signal=48% |
| BP | GO:0042274 | ribosomal small subunit biogenesis | 44 | 0.66634182 | 2.312754094 | 4.01E-07 | 5.17E-05 | 4.88E-05 | 2799 | tags=80%. list=25%. signal=60% |
| BP | GO:0016071 | mRNA metabolic process | 297 | 0.376757729 | 1.779548203 | 8.67E-07 | 0.00010577 | 9.99E-05 | 4157 | tags=60%. list=38%. signal=38% |
| BP | GO:0000375 | RNA splicing. via transesterification reactions | 185 | 0.424369396 | 1.895740584 | 1.53E-06 | 0.00016132 | 0.000152385 | 4157 | tags=65%. list=38%. signal=41% |
| BP | GO:0000377 | RNA splicing. via transesterification reactions with bulged adenosine as nucleophile | 185 | 0.424369396 | 1.895740584 | 1.53E-06 | 0.00016132 | 0.000152385 | 4157 | tags=65%. list=38%. signal=41% |
| BP | GO:0000398 | mRNA splicing. via spliceosome | 185 | 0.424369396 | 1.895740584 | 1.53E-06 | 0.00016132 | 0.000152385 | 4157 | tags=65%. list=38%. signal=41% |
| BP | GO:0008380 | RNA splicing | 199 | 0.415507059 | 1.86923605 | 1.94E-06 | 0.00019528 | 0.000184462 | 4162 | tags=63%. list=38%. signal=40% |
| BP | GO:0009451 | RNA modification | 83 | 0.531933691 | 2.090916639 | 3.21E-06 | 0.00031047 | 0.000293269 | 3723 | tags=73%. list=34%. signal=49% |
| BP | GO:0097485 | neuron projection guidance | 173 | -0.47079489 | -1.860480871 | 4.97E-06 | 0.00045411 | 0.000428956 | 1912 | tags=31%. list=17%. signal=26% |
| BP | GO:0071826 | ribonucleoprotein complex subunit organization | 85 | 0.51289674 | 2.024531276 | 5.09E-06 | 0.00045411 | 0.000428956 | 4145 | tags=68%. list=38%. signal=43% |
| BP | GO:0007411 | axon guidance | 166 | -0.47500683 | -1.869798675 | 5.74E-06 | 0.00049266 | 0.000465367 | 1912 | tags=32%. list=17%. signal=27% |
| BP | GO:0042330 | taxis | 205 | -0.44506765 | -1.791904301 | 7.04E-06 | 0.00058342 | 0.000551097 | 2251 | tags=34%. list=20%. signal=28% |
| BP | GO:0006935 | chemotaxis | 179 | -0.4723648 | -1.875776594 | 7.45E-06 | 0.00059611 | 0.000563084 | 2251 | tags=35%. list=20%. signal=28% |
| BP | GO:0030490 | maturation of SSU-rRNA | 34 | 0.679617898 | 2.229415448 | 8.57E-06 | 0.00066224 | 0.000625552 | 3003 | tags=85%. list=27%. signal=62% |

| | | | | | | | | | | |
|---|---|---|---|---|---|---|---|---|---|---|
| BP | GO:0040011 | locomotion | 399 | -0.38591715 | -1.656846251 | 9.26E-06 | 0.00069298 | 0.000654588 | 2269 | tags=28%. list=21%. signal=23% |
| BP | GO:0022618 | ribonucleoprotein complex assembly | 82 | 0.510650641 | 2.002299753 | 1.27E-05 | 0.0009195 | 0.000868562 | 4145 | tags=68%. list=38%. signal=43% |
| BP | GO:0035295 | tube development | 459 | -0.36713546 | -1.593429986 | 1.31E-05 | 0.00092314 | 0.000871997 | 2352 | tags=28%. list=21%. signal=23% |
| BP | GO:0042273 | ribosomal large subunit biogenesis | 46 | 0.615159436 | 2.15744767 | 1.69E-05 | 0.00115326 | 0.001089372 | 2956 | tags=72%. list=27%. signal=53% |
| BP | GO:0008033 | tRNA processing | 67 | 0.535567368 | 2.028417215 | 1.85E-05 | 0.00122837 | 0.001160314 | 3723 | tags=76%. list=34%. signal=51% |
| BP | GO:0007409 | axonogenesis | 212 | -0.43482508 | -1.756627733 | 2.24E-05 | 0.00144256 | 0.001362645 | 1912 | tags=29%. list=17%. signal=24% |
| BP | GO:0061564 | axon development | 219 | -0.43259222 | -1.755719167 | 3.28E-05 | 0.00205511 | 0.001941252 | 1912 | tags=35%. list=17%. signal=30% |
| BP | GO:0048729 | tissue morphogenesis | 417 | -0.36443661 | -1.568990979 | 5.61E-05 | 0.003421 | 0.003231475 | 2343 | tags=27%. list=21%. signal=22% |
| BP | GO:0032259 | methylation | 102 | 0.460699699 | 1.879285235 | 6.02E-05 | 0.00357764 | 0.003379433 | 3684 | tags=58%. list=33%. signal=39% |
| BP | GO:0045333 | cellular respiration | 64 | 0.520908184 | 1.954734679 | 6.44E-05 | 0.00373643 | 0.003529432 | 3146 | tags=52%. list=29%. signal=37% |
| BP | GO:0035107 | appendage morphogenesis | 231 | -0.39974533 | -1.627623698 | 7.31E-05 | 0.00413479 | 0.003905721 | 2343 | tags=29%. list=21%. signal=24% |
| BP | GO:0035114 | imaginal disc-derived appendage morphogenesis | 230 | -0.4028303 | -1.640558454 | 9.22E-05 | 0.00509186 | 0.004809766 | 2343 | tags=30%. list=21%. signal=24% |
| BP | GO:0006928 | movement of cell or subcellular component | 443 | -0.35671204 | -1.544143107 | 9.68E-05 | 0.00521778 | 0.004928714 | 2269 | tags=26%. list=21%. signal=22% |
| BP | GO:0006400 | tRNA modification | 46 | 0.580528934 | 2.035993796 | 0.000104981 | 0.00553297 | 0.005226442 | 3573 | tags=80%. list=32%. signal=55% |
| BP | GO:0043414 | macromolecule methylation | 99 | 0.460916945 | 1.871771282 | 0.000119152 | 0.00614029 | 0.005800114 | 3524 | tags=56%. list=32%. signal=38% |
| BP | GO:0035120 | post-embryonic appendage morphogenesis | 227 | -0.4009618 | -1.630627247 | 0.000123545 | 0.00622826 | 0.005883209 | 2343 | tags=30%. list=21%. signal=24% |
| BP | GO:0007444 | imaginal disc development | 368 | -0.36516819 | -1.557857785 | 0.000147259 | 0.00702469 | 0.006635516 | 2343 | tags=28%. list=21%. signal=23% |
| BP | GO:0009060 | aerobic respiration | 55 | 0.54262731 | 1.967585947 | 0.000147554 | 0.00702469 | 0.006635516 | 2689 | tags=45%. list=24%. signal=35% |
| BP | GO:0035239 | tube morphogenesis | 346 | -0.36881514 | -1.564855343 | 0.00014843 | 0.00702469 | 0.006635516 | 2468 | tags=30%. list=22%. signal=24% |
| BP | GO:0120036 | plasma membrane bounded cell projection organization | 430 | -0.35362257 | -1.527503559 | 0.000181368 | 0.00841184 | 0.007945824 | 2531 | tags=30%. list=23%. signal=24% |
| BP | GO:0031175 | neuron projection development | 310 | -0.37360465 | -1.570530134 | 0.00023054 | 0.01048279 | 0.009902039 | 2251 | tags=28%. list=20%. signal=23% |
| BP | GO:0048667 | cell morphogenesis involved in neuron differentiation | 283 | -0.38160423 | -1.587834661 | 0.000240152 | 0.01062588 | 0.010037199 | 2251 | tags=28%. list=20%. signal=23% |
| BP | GO:0060562 | epithelial tube morphogenesis | 315 | -0.36938957 | -1.554567339 | 0.000247389 | 0.01062588 | 0.010037199 | 2343 | tags=29%. list=21%. signal=23% |
| BP | GO:0048736 | appendage development | 233 | -0.39413066 | -1.605839831 | 0.000247433 | 0.01062588 | 0.010037199 | 2343 | tags=29%. list=21%. signal=23% |
| BP | GO:0007005 | mitochondrion organization | 145 | 0.3914849 | 1.684410052 | 0.000262414 | 0.01106434 | 0.010451366 | 3796 | tags=55%. list=34%. signal=37% |

| | | | | | | | | | | |
|---|---|---|---|---|---|---|---|---|---|---|
| BP | GO:0008045 | motor neuron axon guidance | 40 | -0.62906453 | -1.971669297 | 0.000280052 | 0.01159363 | 0.010951339 | 1488 | tags=42%. list=13%. signal=37% |
| BP | GO:0001510 | RNA methylation | 44 | 0.552945034 | 1.919173992 | 0.000284966 | 0.01159363 | 0.010951339 | 4030 | tags=80%. list=37%. signal=51% |
| BP | GO:0030182 | neuron differentiation | 494 | -0.33603613 | -1.464792699 | 0.00033167 | 0.01310968 | 0.0123834 | 2303 | tags=26%. list=21%. signal=21% |
| BP | GO:0000462 | maturation of SSU-rRNA from tricistronic rRNA transcript (SSU-rRNA. 5.8S rRNA. LSU-rRNA) | 23 | 0.68736705 | 2.067600762 | 0.000333537 | 0.01310968 | 0.0123834 | 3003 | tags=91%. list=27%. signal=67% |
| BP | GO:0042255 | ribosome assembly | 31 | 0.621329207 | 1.998670829 | 0.000352097 | 0.01360855 | 0.012854634 | 2956 | tags=71%. list=27%. signal=52% |
| BP | GO:0002009 | morphogenesis of an epithelium | 409 | -0.35013134 | -1.506092833 | 0.000362651 | 0.01378667 | 0.01302288 | 2343 | tags=26%. list=21%. signal=22% |
| BP | GO:0007560 | imaginal disc morphogenesis | 273 | -0.38085798 | -1.578233833 | 0.000380947 | 0.01402246 | 0.013245613 | 2343 | tags=29%. list=21%. signal=24% |
| BP | GO:0048563 | post-embryonic animal organ morphogenesis | 273 | -0.38085798 | -1.578233833 | 0.000380947 | 0.01402246 | 0.013245613 | 2343 | tags=29%. list=21%. signal=24% |
| BP | GO:0048737 | imaginal disc-derived appendage development | 232 | -0.39715974 | -1.617859393 | 0.000396063 | 0.01435108 | 0.013556024 | 2343 | tags=29%. list=21%. signal=24% |
| BP | GO:0030030 | cell projection organization | 438 | -0.35133671 | -1.51906468 | 0.000445321 | 0.01588767 | 0.015007487 | 2531 | tags=30%. list=23%. signal=24% |
| BP | GO:0048858 | cell projection morphogenesis | 291 | -0.37541946 | -1.568036291 | 0.00047903 | 0.01658016 | 0.015661615 | 2251 | tags=28%. list=20%. signal=23% |
| BP | GO:0120039 | plasma membrane bounded cell projection morphogenesis | 291 | -0.37541946 | -1.568036291 | 0.00047903 | 0.01658016 | 0.015661615 | 2251 | tags=28%. list=20%. signal=23% |
| BP | GO:0048812 | neuron projection morphogenesis | 289 | -0.37320154 | -1.558200196 | 0.00059345 | 0.02023838 | 0.019117165 | 2251 | tags=28%. list=20%. signal=23% |
| BP | GO:0006626 | protein targeting to mitochondrion | 25 | 0.636309278 | 1.946971104 | 0.000605213 | 0.02034042 | 0.019213553 | 3569 | tags=84%. list=32%. signal=57% |
| BP | GO:0032990 | cell part morphogenesis | 298 | -0.37081852 | -1.552828382 | 0.000626836 | 0.02076617 | 0.019615715 | 1871 | tags=28%. list=17%. signal=24% |
| BP | GO:0048569 | post-embryonic animal organ development | 320 | -0.35861312 | -1.512196425 | 0.000642896 | 0.02099825 | 0.019834933 | 2343 | tags=28%. list=21%. signal=22% |
| BP | GO:0033108 | mitochondrial respiratory chain complex assembly | 44 | 0.532033598 | 1.846594114 | 0.000678889 | 0.0218659 | 0.020654517 | 3541 | tags=70%. list=32%. signal=48% |
| BP | GO:0000904 | cell morphogenesis involved in differentiation | 313 | -0.36784655 | -1.547750831 | 0.00079417 | 0.02494527 | 0.023563293 | 2251 | tags=27%. list=20%. signal=23% |
| BP | GO:0070585 | protein localization to mitochondrion | 30 | 0.617685357 | 1.973996216 | 0.000806768 | 0.02494527 | 0.023563293 | 3316 | tags=77%. list=30%. signal=54% |
| BP | GO:0072655 | establishment of protein localization to mitochondrion | 30 | 0.617685357 | 1.973996216 | 0.000806768 | 0.02494527 | 0.023563293 | 3316 | tags=77%. list=30%. signal=54% |
| BP | GO:0048666 | neuron development | 394 | -0.34479068 | -1.479123615 | 0.000863612 | 0.02635152 | 0.024891636 | 2269 | tags=26%. list=21%. signal=21% |

| | | | | | | | | | | |
|---|---|---|---|---|---|---|---|---|---|---|
| BP | GO:0010257 | NADH dehydrogenase complex assembly | 27 | 0.631255314 | 1.962475038 | 0.000969298 | 0.02881798 | 0.027221451 | 2295 | tags=63%. list=21%. signal=50% |
| BP | GO:0032981 | mitochondrial respiratory chain complex I assembly | 27 | 0.631255314 | 1.962475038 | 0.000969298 | 0.02881798 | 0.027221451 | 2295 | tags=63%. list=21%. signal=50% |
| BP | GO:0000902 | cell morphogenesis | 374 | -0.34045613 | -1.455726961 | 0.001088081 | 0.03193999 | 0.030170504 | 2251 | tags=28%. list=20%. signal=23% |
| BP | GO:0007476 | imaginal disc-derived wing morphogenesis | 201 | -0.37914872 | -1.523574002 | 0.00114329 | 0.03314111 | 0.031305075 | 2343 | tags=28%. list=21%. signal=22% |
| BP | GO:0015980 | energy derivation by oxidation of organic compounds | 94 | 0.413361899 | 1.662721481 | 0.001357367 | 0.03886092 | 0.036708002 | 3316 | tags=45%. list=30%. signal=32% |
| BP | GO:0048707 | instar larval or pupal morphogenesis | 311 | -0.35519737 | -1.493614445 | 0.001435419 | 0.04059434 | 0.038345399 | 2343 | tags=27%. list=21%. signal=22% |
| BP | GO:0007552 | metamorphosis | 322 | -0.34882499 | -1.471196202 | 0.001565016 | 0.04372618 | 0.041303728 | 2343 | tags=27%. list=21%. signal=22% |
| BP | GO:0009887 | animal organ morphogenesis | 490 | -0.32438349 | -1.413273048 | 0.001620565 | 0.04473917 | 0.042260595 | 2343 | tags=26%. list=21%. signal=21% |
| BP | GO:0009886 | post-embryonic animal morphogenesis | 320 | -0.34800009 | -1.46744351 | 0.001658049 | 0.04523547 | 0.042729402 | 2343 | tags=26%. list=21%. signal=21% |
| BP | GO:0018205 | peptidyl-lysine modification | 123 | 0.380342046 | 1.596061415 | 0.001837432 | 0.04954658 | 0.046801673 | 4465 | tags=55%. list=40%. signal=33% |
| CC | GO:0005730 | nucleolus | 156 | 0.597406698 | 2.61231084 | 1.00E-10 | 7.50E-09 | 6.04E-09 | 3092 | tags=70%. list=28%. signal=51% |
| CC | GO:0005840 | ribosome | 129 | 0.610073798 | 2.594172864 | 1.00E-10 | 7.50E-09 | 6.04E-09 | 3136 | tags=73%. list=28%. signal=53% |
| CC | GO:0044391 | ribosomal subunit | 125 | 0.614841844 | 2.601631011 | 1.00E-10 | 7.50E-09 | 6.04E-09 | 3136 | tags=74%. list=28%. signal=53% |
| CC | GO:0098798 | mitochondrial protein-containing complex | 149 | 0.587989897 | 2.553512004 | 1.00E-10 | 7.50E-09 | 6.04E-09 | 3668 | tags=74%. list=33%. signal=50% |
| CC | GO:1990904 | ribonucleoprotein complex | 418 | 0.524281179 | 2.563594817 | 1.00E-10 | 7.50E-09 | 6.04E-09 | 3554 | tags=63%. list=32%. signal=44% |
| CC | GO:0015934 | large ribosomal subunit | 79 | 0.641967298 | 2.511053908 | 2.41E-10 | 1.50E-08 | 1.21E-08 | 2986 | tags=75%. list=27%. signal=55% |
| CC | GO:0005681 | spliceosomal complex | 151 | 0.505746508 | 2.199532354 | 2.05E-08 | 1.10E-06 | 8.86E-07 | 4268 | tags=69%. list=39%. signal=43% |
| CC | GO:0030684 | preribosome | 57 | 0.655981826 | 2.39608271 | 4.75E-08 | 2.09E-06 | 1.68E-06 | 3092 | tags=81%. list=28%. signal=58% |
| CC | GO:0000313 | organellar ribosome | 61 | 0.646519976 | 2.400692094 | 5.65E-08 | 2.09E-06 | 1.68E-06 | 3040 | tags=75%. list=28%. signal=55% |
| CC | GO:0005761 | mitochondrial ribosome | 61 | 0.646519976 | 2.400692094 | 5.65E-08 | 2.09E-06 | 1.68E-06 | 3040 | tags=75%. list=28%. signal=55% |
| CC | GO:0005759 | mitochondrial matrix | 105 | 0.543076951 | 2.234484008 | 6.13E-08 | 2.09E-06 | 1.68E-06 | 3104 | tags=65%. list=28%. signal=47% |
| CC | GO:0022626 | cytosolic ribosome | 68 | 0.612332469 | 2.325191698 | 1.06E-07 | 3.30E-06 | 2.66E-06 | 3783 | tags=85%. list=34%. signal=56% |
| CC | GO:0000315 | organellar large ribosomal subunit | 39 | 0.685676683 | 2.321897784 | 1.23E-06 | 3.28E-05 | 2.64E-05 | 2503 | tags=72%. list=23%. signal=56% |
| CC | GO:0005762 | mitochondrial large ribosomal subunit | 39 | 0.685676683 | 2.321897784 | 1.23E-06 | 3.28E-05 | 2.64E-05 | 2503 | tags=72%. list=23%. signal=56% |

| | | | | | | | | | | |
|---|---|---|---|---|---|---|---|---|---|---|
| CC | GO:0005576 | extracellular region | 181 | -0.48129716 | -1.919148432 | 1.35E-06 | 3.37E-05 | 2.71E-05 | 2144 | tags=41%. list=19%. signal=33% |
| CC | GO:0071013 | catalytic step 2 spliceosome | 101 | 0.500336909 | 2.043321819 | 1.73E-06 | 4.04E-05 | 3.26E-05 | 4677 | tags=80%. list=42%. signal=47% |
| CC | GO:0098800 | inner mitochondrial membrane protein complex | 70 | 0.578631832 | 2.207504898 | 1.83E-06 | 4.04E-05 | 3.26E-05 | 3638 | tags=71%. list=33%. signal=48% |
| CC | GO:0071011 | precatalytic spliceosome | 110 | 0.505195514 | 2.092733235 | 2.04E-06 | 4.25E-05 | 3.42E-05 | 4313 | tags=72%. list=39%. signal=44% |
| CC | GO:0030054 | cell junction | 272 | -0.40474191 | -1.686283619 | 2.43E-05 | 0.0004794 | 0.000386215 | 2118 | tags=30%. list=19%. signal=25% |
| CC | GO:0015935 | small ribosomal subunit | 49 | 0.588289586 | 2.085929342 | 4.91E-05 | 0.00091981 | 0.000741012 | 4005 | tags=86%. list=36%. signal=55% |
| CC | GO:0032040 | small-subunit processome | 29 | 0.690946199 | 2.192091718 | 5.66E-05 | 0.0010109 | 0.000814398 | 3092 | tags=90%. list=28%. signal=65% |
| CC | GO:0005746 | mitochondrial respirasome | 44 | 0.583948974 | 2.021338388 | 0.000118641 | 0.00202229 | 0.001629189 | 3638 | tags=73%. list=33%. signal=49% |
| CC | GO:0005740 | mitochondrial envelope | 167 | 0.393685775 | 1.737640089 | 0.000163087 | 0.00265902 | 0.002142147 | 3667 | tags=58%. list=33%. signal=39% |
| CC | GO:0019866 | organelle inner membrane | 123 | 0.429178472 | 1.811748692 | 0.000191572 | 0.0029298 | 0.002360288 | 3667 | tags=63%. list=33%. signal=42% |
| CC | GO:0022625 | cytosolic large ribosomal subunit | 41 | 0.618974694 | 2.115304194 | 0.000200664 | 0.0029298 | 0.002360288 | 3539 | tags=83%. list=32%. signal=57% |
| CC | GO:0031966 | mitochondrial membrane | 153 | 0.40187827 | 1.754324724 | 0.000206875 | 0.0029298 | 0.002360288 | 3667 | tags=58%. list=33%. signal=39% |
| CC | GO:0098803 | respiratory chain complex | 43 | 0.568899863 | 1.959205478 | 0.000210946 | 0.0029298 | 0.002360288 | 3638 | tags=70%. list=33%. signal=47% |
| CC | GO:0005743 | mitochondrial inner membrane | 116 | 0.424995959 | 1.777658461 | 0.000235115 | 0.00314886 | 0.002536766 | 3238 | tags=56%. list=29%. signal=40% |
| CC | GO:0005615 | extracellular space | 110 | -0.48079537 | -1.801148122 | 0.000286093 | 0.00369948 | 0.002980355 | 2457 | tags=43%. list=22%. signal=34% |
| CC | GO:0030532 | small nuclear ribonucleoprotein complex | 47 | 0.556474577 | 1.955408 | 0.000317674 | 0.00397093 | 0.003199037 | 4728 | tags=85%. list=43%. signal=49% |
| CC | GO:0120114 | Sm-like protein family complex | 48 | 0.557265236 | 1.968295782 | 0.000359581 | 0.00408737 | 0.003292845 | 4728 | tags=85%. list=43%. signal=49% |
| CC | GO:0030686 | 90S preribosome | 22 | 0.693133474 | 2.05249486 | 0.000399499 | 0.00408737 | 0.003292845 | 3003 | tags=86%. list=27%. signal=63% |
| CC | GO:0005747 | mitochondrial respiratory chain complex I | 28 | 0.650271867 | 2.040028787 | 0.000402598 | 0.00408737 | 0.003292845 | 2372 | tags=68%. list=21%. signal=53% |
| CC | GO:0030964 | NADH dehydrogenase complex | 28 | 0.650271867 | 2.040028787 | 0.000402598 | 0.00408737 | 0.003292845 | 2372 | tags=68%. list=21%. signal=53% |
| CC | GO:0045271 | respiratory chain complex I | 28 | 0.650271867 | 2.040028787 | 0.000402598 | 0.00408737 | 0.003292845 | 2372 | tags=68%. list=21%. signal=53% |
| CC | GO:0031967 | organelle envelope | 257 | 0.336066265 | 1.567665514 | 0.000403287 | 0.00408737 | 0.003292845 | 3572 | tags=48%. list=32%. signal=33% |
| CC | GO:0031975 | envelope | 257 | 0.336066265 | 1.567665514 | 0.000403287 | 0.00408737 | 0.003292845 | 3572 | tags=48%. list=32%. signal=33% |
| CC | GO:0070469 | respirasome | 45 | 0.575541804 | 2.00524766 | 0.000429728 | 0.00424074 | 0.003416396 | 3146 | tags=64%. list=29%. signal=46% |
| CC | GO:0070161 | anchoring junction | 103 | -0.4752911 | -1.764260002 | 0.00047976 | 0.00461308 | 0.003716363 | 2409 | tags=39%. list=22%. signal=31% |
| CC | GO:0062023 | collagen-containing extracellular matrix | 15 | -0.74079923 | -1.869053927 | 0.000805233 | 0.00754906 | 0.006081625 | 1648 | tags=67%. list=15%. signal=57% |

| | | | | | | | | | | |
|---|---|---|---|---|---|---|---|---|---|---|
| CC | GO:0045177 | apical part of cell | 114 | -0.45216974 | -1.700156105 | 0.001179907 | 0.01079183 | 0.008694053 | 1967 | tags=32%. list=18%. signal=26% |
| CC | GO:0030424 | axon | 128 | -0.43850885 | -1.674170882 | 0.001237806 | 0.01105184 | 0.008903518 | 2852 | tags=41%. list=26%. signal=31% |
| CC | GO:0097525 | spliceosomal snRNP complex | 38 | 0.5638852 | 1.895775425 | 0.001692752 | 0.01476237 | 0.011892771 | 4728 | tags=89%. list=43%. signal=51% |
| CC | GO:1990204 | oxidoreductase complex | 51 | 0.501363551 | 1.794040386 | 0.001845012 | 0.01572453 | 0.012667905 | 3146 | tags=53%. list=29%. signal=38% |
| CC | GO:0030880 | RNA polymerase complex | 67 | 0.457006478 | 1.728292544 | 0.002117641 | 0.01764701 | 0.014216676 | 3913 | tags=60%. list=35%. signal=39% |
| CC | GO:0055029 | nuclear DNA-directed RNA polymerase complex | 65 | 0.456628069 | 1.715932597 | 0.002198242 | 0.01792045 | 0.01443697 | 3913 | tags=60%. list=35%. signal=39% |
| CC | GO:0000428 | DNA-directed RNA polymerase complex | 66 | 0.462411508 | 1.742115359 | 0.003185962 | 0.02541991 | 0.020478636 | 3913 | tags=61%. list=35%. signal=39% |
| CC | GO:0043005 | neuron projection | 189 | -0.37439425 | -1.500580222 | 0.004025765 | 0.03129346 | 0.025210453 | 2535 | tags=33%. list=23%. signal=26% |
| CC | GO:0022627 | cytosolic small ribosomal subunit | 28 | 0.595505142 | 1.868214962 | 0.004137337 | 0.03129346 | 0.025210453 | 3783 | tags=82%. list=34%. signal=54% |
| CC | GO:0005732 | sno(s)RNA-containing ribonucleoprotein complex | 15 | 0.694516451 | 1.865097983 | 0.004172462 | 0.03129346 | 0.025210453 | 3092 | tags=87%. list=28%. signal=62% |
| CC | GO:0000314 | organellar small ribosomal subunit | 22 | 0.602358116 | 1.78369244 | 0.004617254 | 0.0332975 | 0.026824937 | 4005 | tags=91%. list=36%. signal=58% |
| CC | GO:0005763 | mitochondrial small ribosomal subunit | 22 | 0.602358116 | 1.78369244 | 0.004617254 | 0.0332975 | 0.026824937 | 4005 | tags=91%. list=36%. signal=58% |
| CC | GO:0098573 | intrinsic component of mitochondrial membrane | 32 | 0.52420858 | 1.702976313 | 0.006240639 | 0.04415546 | 0.035572262 | 4095 | tags=72%. list=37%. signal=45% |
| CC | GO:0030688 | preribosome. small subunit precursor | 10 | 0.739953136 | 1.783423984 | 0.007165249 | 0.04975867 | 0.040086283 | 2289 | tags=80%. list=21%. signal=63% |

**Table S6. Enrichment analysis**. Summary of the results

| **Enriched pthways/GO (padj<0.05)** | **Total** | **Positive** | **Negative** |
|---|---|---|---|
| KEGG | 26 | 10 | 16 |
| BP | 86 | 46 | 40 |
| MF | 17 | 12 | 5 |
| CC | 54 | 46 | 8 |

**Table S7. Sources of data for those species for which E93 expression in embryos and pre-adult stage was obtained.**

Data from genomes and transcriptomes

| N | E93 orthologue | Species | Metamorphosis | Assembly name | Assembly accession | Number of mapped SRA libraries | Number of analyzed SRA libraries | Comment |
|---|---|---|---|---|---|---|---|---|
| 1 | E93 | *Blattella germanica* | Hemimetabolan | x | x | 14 | 12 | DOI: 10.1038/s42003-021-02197-9 |
| 2 | E93 | *Ischnura senegalensis* | Hemimetabolan | x | x | x | 33 | DOI:10.1073/pnas.2114773119 |
| 3 | GBI_04085 | *Gryllus bimaculatus* | Hemimetabolan | x | x | 51 | 34 | DOI: 10.1016/bs.ctdb.2022.02.003 |
| 4 | gene-LOC110851286 | *Folsomia candida* | Ametabolan | ASM221717v1 | GCF_002217175.1 | 11 | 11 | BGI-seq |
| 5 | gene-LOC100161761 | *Acyrthosiphon pisum* | Hemimetabolan | pea_aphid_22Mar2018_4r6ur | GCA_005508785.1 | 12 | 11 | |
| 6 | gene-LOC111052728 | *Nilaparvata lugens* | Hemimetabolan | ASM1435652v1 | GCA_014356525.1 | 10 | 10 | |
| 7 | gene-LOC114128046 | *Aphis gossypii* | Hemimetabolan | ASM401081v1 | GCA_004010815.1 | 6 | 6 | Embryo: Illumina. Post-embryo: BGI-seq. |
| 8 | gene-CLODIP_2_CD15741 | *Cloeon dipterum* | Hemimetabolan | CLODIP2 | GCA_902829235.1 | 31 | 31 | |
| 9 | gene-LOC110827285 | *Zootermopsis nevadensis* | Hemimetabolan | ZooNev1.0 | GCF_000696155.1 | 3 | 3 | |
| 10 | gene-LOC109033008 | *Bemisia tabaci* | Hemimetabolan | ASM185493v1 | GCF_001854935.1 | 29 | 26 | |
| 11 | gene-LOC103516559 | *Diaphorina citri* | Hemimetabolan | Diaci psyllid genome assembly version 1.1 | GCF_000475195.1 | 11 | 11 | |
| 12 | gene-LOC6046300 | *Culex quinquefasciatus* | Holometabolan | VPISU_Cqui_1.0_pri_paternal | GCA_015732765.1 | 8 | 7 | |
| 13 | gene-LOC105215468 | *Zeugodacus cucurbitae* | Holometabolan | ASM80634v1 | GCF_000806345.1 | 29 | 29 | |

| | | | | | | | |
|---|---|---|---|---|---|---|---|
| 14 | gene-LOC105255462 | *Camponotus floridanus* | Holometabolan | Cflo_v7.5 | GCA_003227725.1 | 9 | 9 |
| 15 | gene-Mblk-1 | *Apis mellifera* | Holometabolan | Amel_HAv3.1 | GCA_003254395.2 | 90 | 82 |
| 16 | gene-LOC655012 | *Tribolium castaneum* | Holometabolan | Tcas5.2 | GCA_000002335.3 | 85 | 85 |
| 17 | gene-LOC105183699 | *Harpegnathos saltator* | Holometabolan | Hsal_v8.5 | GCA_003227715.1 | 7 | 7 |
| 18 | gene-LOC115453157 | *Manduca sexta* | Holometabolan | JHU_Msex_v1.0 | GCA_014839805.1 | 9 | 9 |
| 19 | gene-LOC105385744 | *Plutella xylostella* | Holometabolan | Haplomerged_assembly | GCF_905116875.1 | 2 | 2 |
| 20 | gene-LOC101737038 | *Bombyx mori* | Holometabolan | Bmori_2016v1.0 | GCF_014905235.1 | 69 | 65 |
| 21 | gene-LOC122503059 | *Leptopilina heterotoma* | Holometabolan | ASM1547642v1 | GCF_015476425.1 | 4 | 4 |
| 22 | gene-LOC116848284 | *Odontomachus brunneus* | Holometabolan | Obru_v1 | GCF_010583005.1 | 4 | 4 |
| 23 | gene-LOC118503280 | *Anopheles stephensi* | Holometabolan | UCI_ANSTEP_V1.0 | GCF_013141755.1 | 4 | 4 |
| 24 | gene-LOC105832848 | *Monomorium pharaonis* | Holometabolan | ASM1337386v2 | GCF_013373865.1 | 15 | 15 |
| 25 | gene-LOC119657514 | *Hermetia illucens* | Holometabolan | iHerIll2.2.curated.20191125 | GCF_905115235.1 | 8 | 8 |
| 26 | gene-LOC106106135 | *Papilio polytes* | Holometabolan | Ppol_1.0 | GCF_000836215.1 | 67 | 67 |
| 27 | gene-LOC105226331 | *Bactrocera dorsalis* | Holometabolan | ASM78921v2 | GCF_000789215.1 | 27 | 25 |
| 28 | gene-LOC113396009 | *Vanessa tameamea* | Holometabolan | ASM293899v1 | GCF_002938995.1 | 2 | 2 |
| 29 | gene-LOC111002879 | *Pieris rapae* | Holometabolan | ilPieRapa1.1 | GCF_905147795.1 | 9 | 8 |
| 30 | gene-HW555_010051 | *Spodoptera exigua* | Holometabolan | NJAU_Sexi_v1 | GCA_011316535.1 | 15 | 15 |
| 31 | Dmel_CG18389 | *Drosophila melanogaster* | Holometabolan | Release 6 plus ISO1 MT | GCA_000001215.4 | 179 | 179 |

| | | | | | | | | |
|---|---|---|---|---|---|---|---|---|
| 32 | E93_homolog | *Trichomalopsis sarcophagae* | Holometabolan | ASM224990v1 | GCA_002249905.1 | 6 | 6 | E93 manually annotated based on blast |
| 33 | NO E93 | *Megalopta genalis* | Holometabolan | USU_MGEN_1.2 | GCF_011865705.1 | 12 | 0 | E93 not found in the genomic seqence |
| 34 | gene-LOC106614581 | *Bactrocera oleae* | Holometabolan | ASM78921v2 | GCF_001188975.3 | 10 | 0 | No E93 expression |

Data from qRT-PCR measurements

| N | E93 orthologue | Species | Metamorphosis | Reference |
|---|---|---|---|---|
| 1 | E93 | *Blattella germanica* | Hemimetabolan | Present work and DOI: 10.1016/j.ibmb.2014.06.009 |
| 2 | E93 | *Frankliniella occidentalis* | Neometabolan | DOI: 10.1371/journal.pone.0254963 |
| 3 | E93 | *Haplothrips brevitubus* | Neometabolan | DOI: 10.1371/journal.pone.0254963 |
| 4 | E93 | *Planococcus kraunhiae* | Neometabolan | DOI: 10.1016/j.ibmb.2018.11.008 |
| 5 | E93 | *Thermobia domestica* | Ametabolan | Present work |

**Table S8. E93 Ratios in the analyzed species.**

| Species | Metamorphosis | Ratio | Log2ratio | Method |
|---|---|---|---|---|
| *Acyrthosiphon pisum* | Neometabolan | 0.872437016 | -0.196877112 | RNA-Seq |
| *Anopheles stephensi* | Holometabolan | 4.213382907 | 2.074979032 | RNA-Seq |
| *Aphis gossypii* | Neometabolan | 2.153841014 | 1.106911761 | RNA-Seq |
| *Apis mellifera* | Holometabolan | 65.80996543 | 6.040234159 | RNA-Seq |
| *Bactrocera dorsalis* | Holometabolan | 2.336939935 | 1.224620654 | RNA-Seq |
| *Bemisia tabaci* | Neometabolan | 8.004720246 | 3.000850983 | RNA-Seq |
| *Blattella germanica* | Hemimetabolan | 1.703673413 | 0.768648803 | RNA-Seq |
| *Bombyx mori* | Holometabolan | 7.890068609 | 2.980037845 | RNA-Seq |
| *Camponotus floridanus* | Holometabolan | 5.7268811 | 2.517749651 | RNA-Seq |
| *Cloeon dipterum* | Hemimetabolan | 2.323999519 | 1.21660977 | RNA-Seq |
| *Culex quinquefasciatus* | Holometabolan | 586.0910119 | 9.194980903 | RNA-Seq |
| *Diaphorina citri* | Neometabolan | 30.60292493 | 4.935597643 | RNA-Seq |
| *Drosophila melanogaster* | Holometabolan | 101.1238707 | 6.659979782 | RNA-Seq |
| *Folsomia candida* | Ametabolan | 0.300142559 | -1.736280192 | RNA-Seq |
| *Gryllus bimaculatus* | Hemimetabolan | 0.22985733 | -2.12118942 | RNA-Seq |
| *Harpegnathos saltator* | Holometabolan | 3.8335012 | 1.938662631 | RNA-Seq |
| *Hermetia illucens* | Holometabolan | 2.564433759 | 1.358640306 | RNA-Seq |
| *Ischnura senegalensis* | Hemimetabolan | 58.6025641 | 5.872891885 | RNA-Seq |
| *Leptopilina heterotoma* | Holometabolan | 11.15337546 | 3.479408489 | RNA-Seq |
| *Manduca sexta* | Holometabolan | 23.54274384 | 4.557210567 | RNA-Seq |
| *Monomorium pharaonis* | Holometabolan | 3.979897094 | 1.992731128 | RNA-Seq |
| *Nilaparvata lugens* | Hemimetabolan | 1.045490367 | 0.06417977 | RNA-Seq |
| *Odontomachus brunneus* | Holometabolan | 9.731631342 | 3.282681668 | RNA-Seq |
| *Papilio polytes* | Holometabolan | 2.535376094 | 1.34219977 | RNA-Seq |
| *Pieris rapae* | Holometabolan | 15.92365664 | 3.993099763 | RNA-Seq |
| *Plutella xylostella* | Holometabolan | 14.36420465 | 3.844406207 | RNA-Seq |
| *Spodoptera exigua* | Holometabolan | 1.892437596 | 0.920245728 | RNA-Seq |
| *Tribolium castaneum* | Holometabolan | 26.5635343 | 4.731375206 | RNA-Seq |
| *Trichomalopsis sarcophagae* | Holometabolan | 38.77087605 | 5.276901429 | RNA-Seq |
| *Vanessa tameamea* | Holometabolan | 1.750162218 | 0.807488648 | RNA-Seq |
| *Zeugodacus cucurbitae* | Holometabolan | 667.0357573 | 9.381620291 | RNA-Seq |
| *Zootermopsis nevadensis* | Hemimetabolan | 3.224058197 | 1.688877786 | RNA-Seq |
| *Blattella germanica* | Hemimetabolan | 1.460176991 | 0.546143252 | qPCR |
| *Frankliniella occidentalis* | Neometabolan | 9.416666667 | 3.235216462 | qPCR |
| *Haplothrips brevitubus* | Neometabolan | 9500 | 13.2137118 | qPCR |
| *Planococcus kraunhiae* | Neometabolan | 95.93276344 | 6.583951711 | qPCR |
| *Thermobia domestica* | Ametabolan | 0.52 | -0.943416472 | qPCR |

**Table S9. Primer sequences used for qRT-PCR and RNAi experiments**

| Gene name | Reference | Forward primer sequence | Reverse primer sequence |
|---|---|---|---|
| ***Blattella germanica*** | | | |
| qRT-PCR | | | |
| *Actin 5c* | Acc. N. AJ862721.1 | AGCTTCCTGATGGTCAGGTGA | TGTCGGCAATTCCAGGGTACATGGT |
| *eve* | Ylla et al., 2018* | AGGTGCGGGATTGTTAACTG | AGGGTTGGAAAAGCTTTGGT |
| *E93* | Acc. N. HF536494 | TCCAATGTTTGATCCTGCAA | TTTGGGATGCAAAGAAATCC |
| *Ftz* | Ylla et al., 2018* | AACGCCTTCTTCGAAGTCAA | TTGAAATTGCTGGTGCCATA |
| *knrl* | Ylla et al., 2018* | ATGCTAGGACACCCTGCTCA | TGACGGAAGGCGATGTTGAGT |
| *Kr* | Acc. N. LT717630 | CGTACACACACGGGAGAAAA | AACAAATTGCCGGTCACAAT |
| *nvd* | Ylla et al., 2018* | CTGGGGCCAGTCACAATACT | GCAGGGGCTTGTCAATGTAT |
| *Ppa* | Ylla et al., 2018* | TAAGTCGGTATGGCGAGGAG | CCACGTCCCTAAGACTACGC |
| *shd* | Ylla et al., 2018* | CACAGAGGCGCACAAGTTTA | GTTCCCCTTCAAAGTCCACA |
| *sina* | Ylla et al., 2018* | ATGTGGCAATAGAGGGCTTG | TACAGAATTTGGAGGGATTA |
| *stv* | Ylla et al., 2018* | AGGAGAAGACGACACGGCTA | TGTTACGAAGCCCATGTTGA |
| *ths* | Ylla et al., 2018* | CGAAAGTGGAAGCTGGTAGG | ATGGTCTTCCTCGACGATTG |
| *Bger_02424* | Ylla et al., 2018* | TGGTCTTCATGCCTCTGTTG | AGACGGCAACTATGGAATCG |
| *Bger_15646* | Ylla et al., 2018* | ATATTGCAGCTGGCGTTCAT | TGAGCCATCAGGAGAGTGAA |
| *Bger_20604* | Ylla et al., 2018* | AACAGCAACCTGTCATGCAA | ATTTCGCAAATGGCTGACTT |
| *Bger_23358* | Ylla et al., 2018* | CGGTGGAAGAGAAGGACTTG | CCTGAGCAACAGAAAGCACA |
| *Bger_27721* | Ylla et al., 2018* | CTCGCAGACGGAGAACAAGT | GTTGTTCCCGGATGTGTCTC |
| dsRNA | | | |
| E93 dsRNA | Acc. N. HF536494 | AAAGAGTTGTCGGGAGCAGA | CCACTGCTAGAAGCCACTCC |
| polyhedrin | Acc. N. K01149 | ATCCTTTCCTGGGACCCGGCA | ATGAAGGCTCGACGATCCTA |
| ***Thermobia domestica*** | | | |
| qRT-PCR | | | |
| E93 | Belles, 2019** | TGGGGCTACGAGTGTGAAAG | CGCAGTGCTGTCATCATTCG |
| Kr-h1 | Acc. N. JN416989.1 | TCGTCTCCGGATCCAGAGAG | TGGACATTGGTACTTCCGGC |
| Br-C | Acc. N. GQ983556.1 | GGGCTCATCAGGGTTTTCCA | GGATTTGATTCGCTGGCGTC |
| rp49/RpL32 | Acc. N. AB689035.1 | GCACCAGAGTGACCGATATGTTAA | CGGACTCTGTTATCAATACCTTTGG |

* Ylla, G., Piulachs, M.D., Belles, X. 2018. Comparative transcriptomics in two extreme neopterans reveals general trends in the evolution of modern insects. *iScience* **4**, 164–179. https://doi.org/10.1016/j.isci.2018.05.017

** Belles, X. 2019. The innovation of the final moult and the origin of insect metamorphosis. *Philos. Trans. R. Soc. B Biol. Sci.* **374**, 20180415. https://doi.org/10.1098/rstb.2018.0415

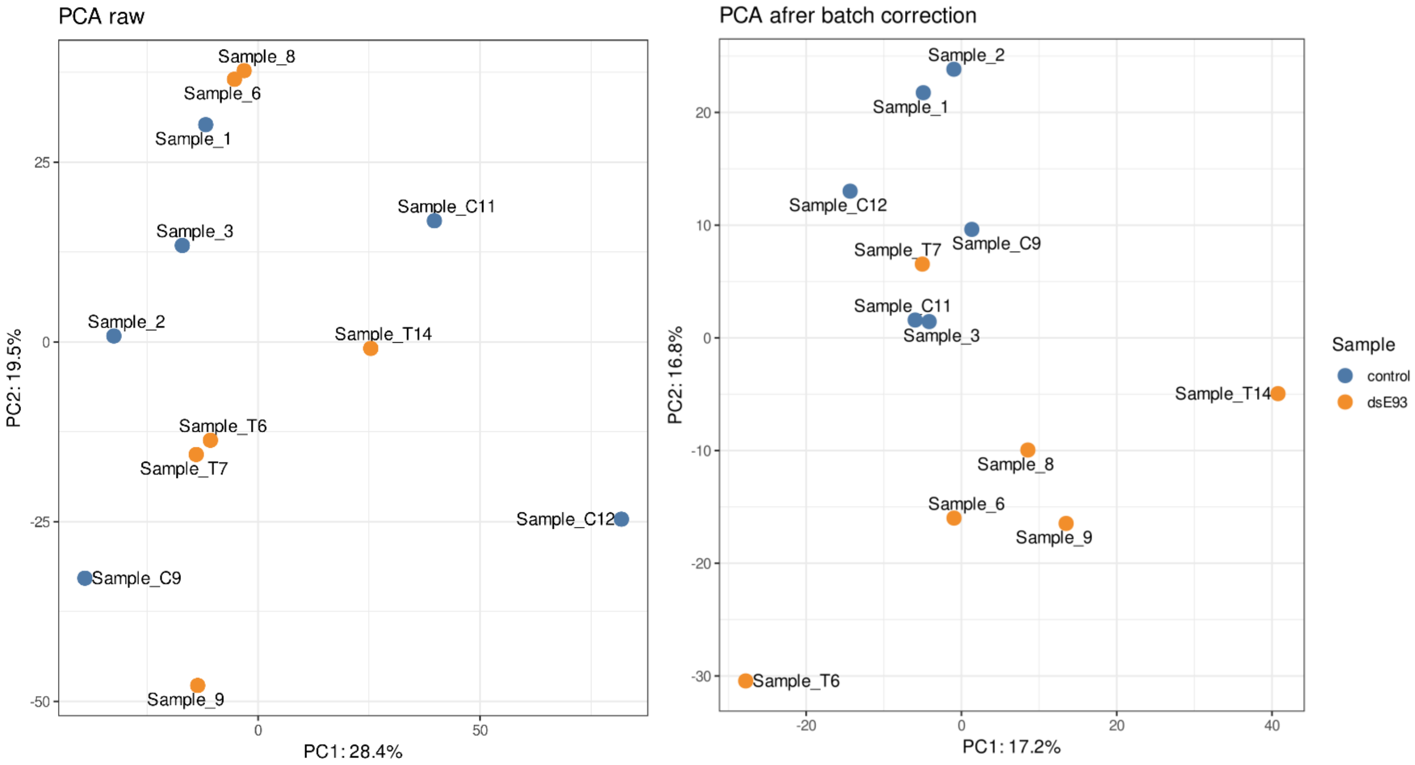

**Figure S1. Principal Component Analysis of all available libraries.** Analysis of the different batches (PCA row), and results after removing the batch effect (PCA after batch correction).

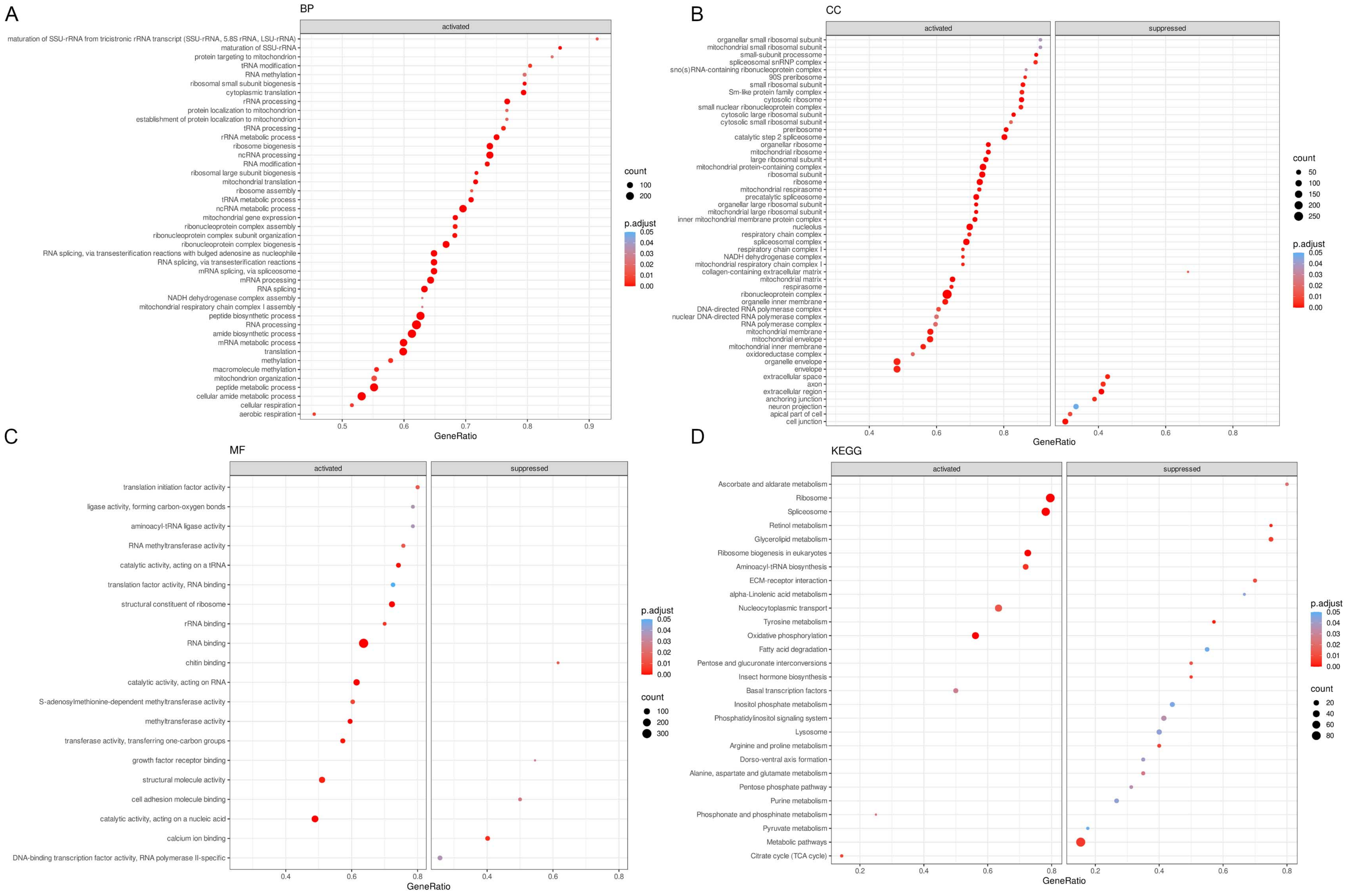

**Figure S2. Gene Set Enrichment Analysis (GSEA) of the RNA-seq from the E93 depletion experiment.** (A-C) Significantly activated GO terms of biological processes (A: BP), cellular components (B: CC) and molecular functions (C: MF); regarding BP, the terms that resulted down-regulated after the E93 depletion are shown in Figure 2. (D) Significant KEGG pathways.

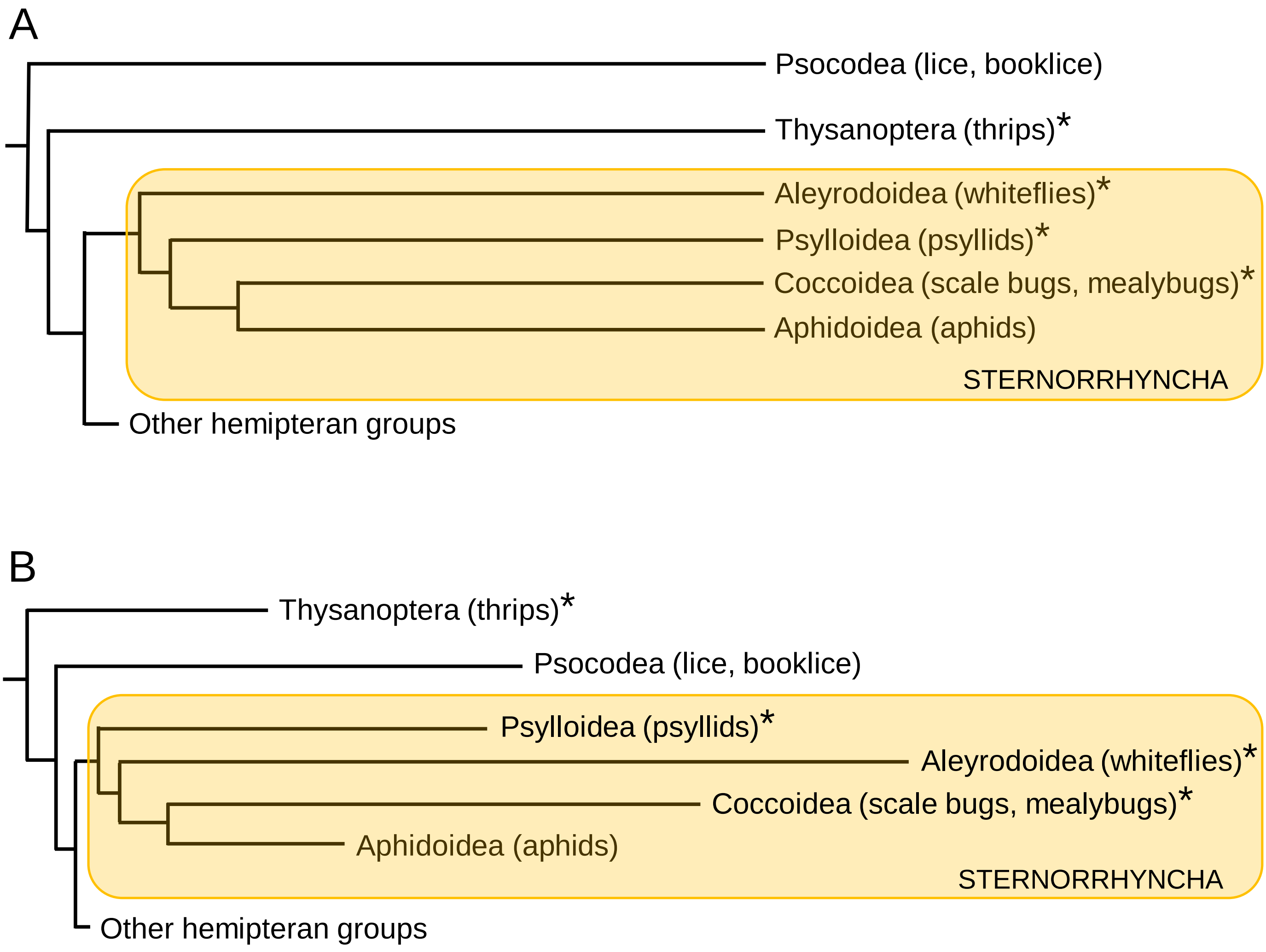

**Figure S3. Phylogeny the Hemiptera Sternorrhyncha, and relationships with Thysanoptera and Psocodea**. (A) According to Johnson et al., 2018, based on maximum likelihood approaches and protein-coding gene sequences. (B). According to Song et al., 2019, based on maximum likelihood approaches and mitochondrial genome data.

* Groups with juvenile stages showing a clearly different morphology from the adult.
